# The *cis*-regulatory dynamics of embryonic development at single cell resolution

**DOI:** 10.1101/166066

**Authors:** Darren A. Cusanovich, James P. Reddington, David A. Garfield, Riza Daza, Raquel Marco-Ferreres, Lena Christiansen, Xiaojie Qiu, Frank Steemers, Cole Trapnell, Jay Shendure, Eileen E.M. Furlong

**Author notes:** These authors contributed equally (Darren A. Cusanovich, James P. Reddington, David A. Garfield).

## Abstract

Single cell measurements of gene expression are providing new insights into lineage commitment, yet the regulatory changes underlying individual cell trajectories remain elusive. Here, we profiled chromatin accessibility in over 20,000 single nuclei across multiple stages of *Drosophila* embryogenesis. Our data reveal heterogeneity in the regulatory landscape prior to gastrulation that reflects anatomical position, a feature that aligns with future cell fate. During mid embryogenesis, tissue granularity emerges such that cell types can be inferred by their chromatin accessibility, while maintaining a signature of their germ layer of origin. We identify over 30,000 distal elements with tissue-specific accessibility. Using transgenic embryos, we tested the germ layer specificity of a subset of predicted enhancers, achieving near-perfect accuracy. Overall, these data demonstrate the power of shotgun single cell profiling of embryos to resolve dynamic changes in open chromatin during development, and to uncover the *cis*-regulatory programs of germ layers and cell types.

## MAIN TEXT

The remarkable power of *Drosophila melanogaster* as a model for studying embryogenesis derives from a sophisticated repertoire of genetic tools and the accumulated depth of knowledge of its development. Nonetheless, our current understanding of how the regulation of embryogenesis is encoded by the genome in *Drosophila*, and other multicellular organisms, stems from an averaged view generated from thousands to millions of cells. The dynamics of the gene regulatory landscape as germ layers and cell types emerge during embryogenesis remains poorly understood, in part because we have achieved only piecemeal characterization of the chromatin landscape with traditional genomic technologies.

To address these challenges, we set out to measure the regulatory landscape of chromatin accessibility during *Drosophila* embryogenesis at single cell resolution. Technologies for profiling various aspects of single cell biology have recently proliferated, but most efforts have focused on single cell RNA sequencing, which quantifies the transcriptional output of the genome. Recently, we (and others) developed methods for quantifying genome-wide chromatin accessibility in single nuclei^1,2^, as chromatin accessibility is a powerful indicator of the regulatory DNA active in different cell types. The method we developed, single cell combinatorial indexing assay for transposase accessible chromatin (sci-ATAC-seq), uses combinatorial indexing to facilitate the profiling of genome-wide chromatin accessibility in large numbers of single nuclei per experiment^1^.

### A single cell map of open chromatin during embryogenesis

We adapted our sci-ATAC-seq protocol to work with nuclei obtained from *Drosophila* embryos, concurrently implementing optimizations to increase the sensitivity of the protocol and enable compatibility with formaldehyde fixation. To study the dynamics of chromatin accessibility during developmental progression, we isolated intact nuclei from fixed embryos spanning three landmark cell states in embryogenesis: 2-4 hours (hrs) after egg laying (predominantly stage 5), when cells are multipotent, including blastoderm nuclei; 6-8 hrs (predominantly stage 11), to capture a midpoint in embryonic development when major lineages in the mesoderm and ectoderm are specified; and 10-12 hrs (predominantly stage 13), when terminal tissue differentiation is occurring (Fig. 1a). The nuclei processed from each time point were derived from thousands of embryos (of both sexes). As embryogenesis is not perfectly synchronous the processed nuclei naturally sample intermediate developmental states, thereby providing a comprehensive range of cellular transitions during embryogenesis.

**Figure 1.**
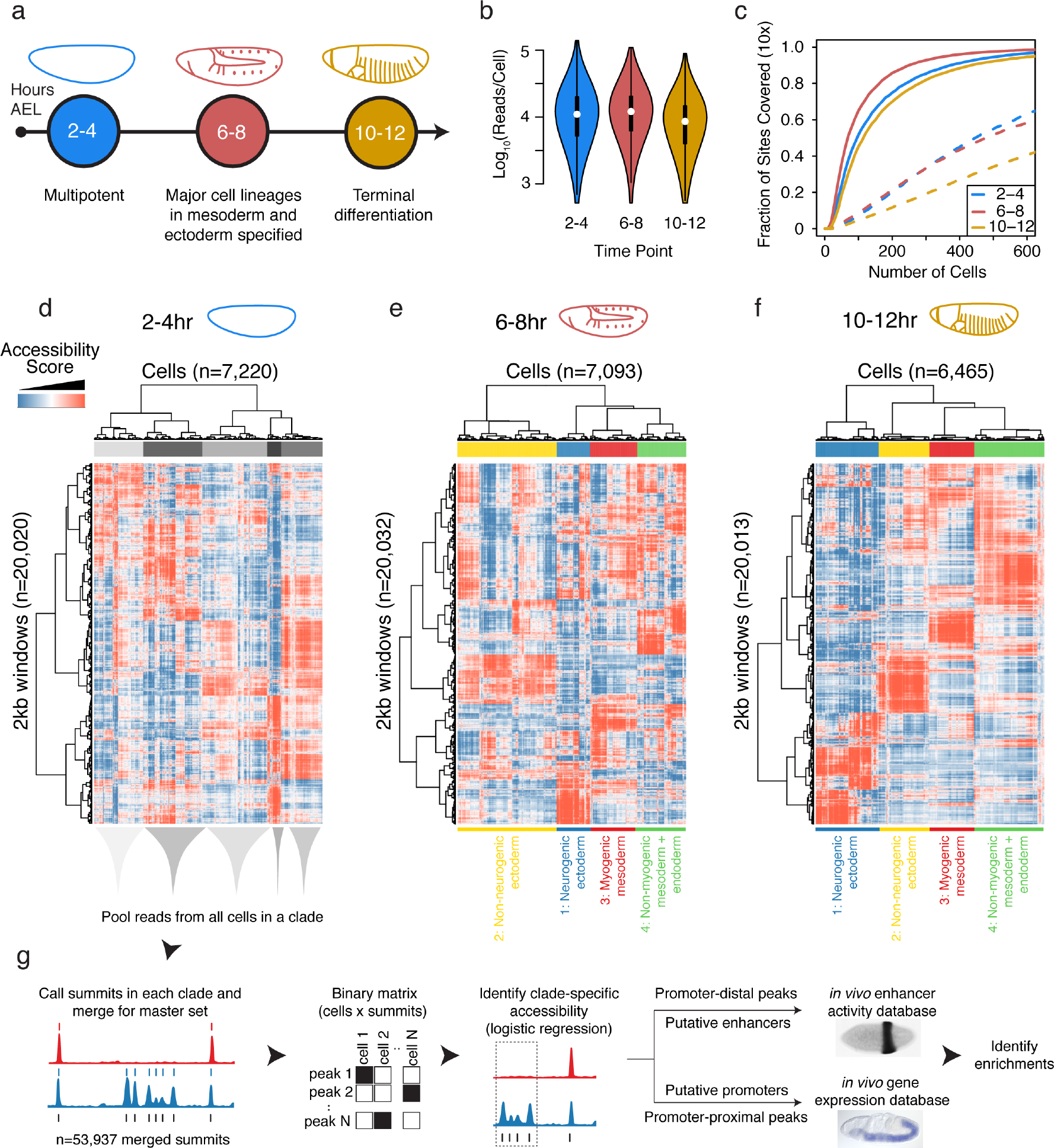
Single cell profiling of chromatin accessibility across *Drosophila* embryogenesis. **a**, Timeline of sampling corresponds to major milestones of embryogenesis (‘AEL’ = after egg laying). **b**, Distribution of unique, mappable reads per cell at each of the three developmental time points. **c**, Fraction of previously characterized DHS covered in least 10 cells upon sampling a given number of cells (solid lines) as compared to random genomic windows (dashed lines). **d-f**, Heatmaps of binarized, LSI-transformed, and clustered read counts in 2 kb windows across the genome at 2-4 hr (d), 4-6 hr (e) and 10-12 hr (f) after egg laying, showing the relationships among single cells (columns) and windows (rows) based on patterns of shared accessibility. Major clades are readily assignable to germ layers at post-gastrulation time points (e,f). g, Approach taken to annotate clades by intersecting clade-specific peaks of chromatin accessibility with enhancer activity and gene expression data. The *in situ* image of enhancer activity (black stain) was obtained from^5^ and the RNA *in situ* (blue stain) was obtained from the Berkeley Drosophila Genome Project^8,30,31^.

Of the 430 million read pairs sequenced in total, 71% mapped to the nuclear reference genome (FlyBase 5; mapping quality ≥10) and had an assigned cell barcode. As observed previously^1^, the logtransformed counts of sci-ATAC-seq reads assigned to each possible cell barcode were bimodally distributed, with peaks corresponding to background noise and successfully indexed cells (Extended Data Fig. 1a). We used a Gaussian mixture model to define the minimum depth for calling a cell barcode as valid at 518 reads (Methods). At this threshold, we sequenced a median of 20,587 reads per cell, of which a median of 10,540 (average 13,349 ± 11,349) per cell were retained after de-duplication (Fig. 1b). Due to our protocol optimizations, this is approximately an order of magnitude more reads per cell than our initial description of sci-ATAC-seq^1^. Reassuringly, the fragment size distribution was consistent with expectation for ATAC-seq data at each time point (Extended Data Fig. 1b). Overall, we recovered genome-wide chromatin accessibility profiles for 23,086 cells across the three time points. Reads were enriched in open chromatin regions as defined by bulk DNase-seq on *Drosophila* embryos; conversely, DNase hypersensitive sites (DHS; an alternative measure of accessible chromatin) were well-covered by our sci-ATAC-seq data (Fig. 1c)^3^.

As a first step to analyze this dataset, we partitioned the genome into 2 kilobase (kb) windows and scored each cell by whether there were any reads observed in each window. As this process results in a sparse, binary matrix of cells (columns) by genomic windows (rows), we retained only the 20,000 most frequently accessible windows, removed the sparsest 10% of cells, and then performed latent semantic indexing (LSI) to reduce the dimensionality of this matrix, as previously described^1^ (Methods).

Biclustering of the LSI matrices for each developmental time point revealed subsets of cells exhibiting similar chromatin accessibility (Fig. 1d-f). To more finely map the regulatory elements underlying this clustering, we aggregated data from all cells within each of the largest four to five clades per time point (Fig. 1g). This ‘*in silico* sorting’ yielded datasets for each clade with sufficient read depth to identify broader peaks of accessibility as well as punctuated summits of accessibility. Merging summits across all three time points and clades, we generated a master set of 53,937 potential *cis*-regulatory elements. To determine which regulatory elements exhibited clade-specific chromatin accessibility, we applied a logistic regression framework (Methods; Table S1), identifying 41,405 sites (77%) with clade-specific accessibility in at least one time point. These include 12,741 sites that were specifically more open in one clade relative to others at the 2-4 hr time point, 25,728 such sites at 6-8 hrs, and 28,614 at 10-12 hrs (Extended Data Fig. 2). These results confirm that the regulatory landscape of the developing embryo is both highly heterogeneous and dynamic. About twice as many clade-specific differentially accessible sites are identified at later time points compared to 2-4 hrs, which likely reflects the increase in complexity and refinement of cell identities from their common progenitors.

**Figure 2.**
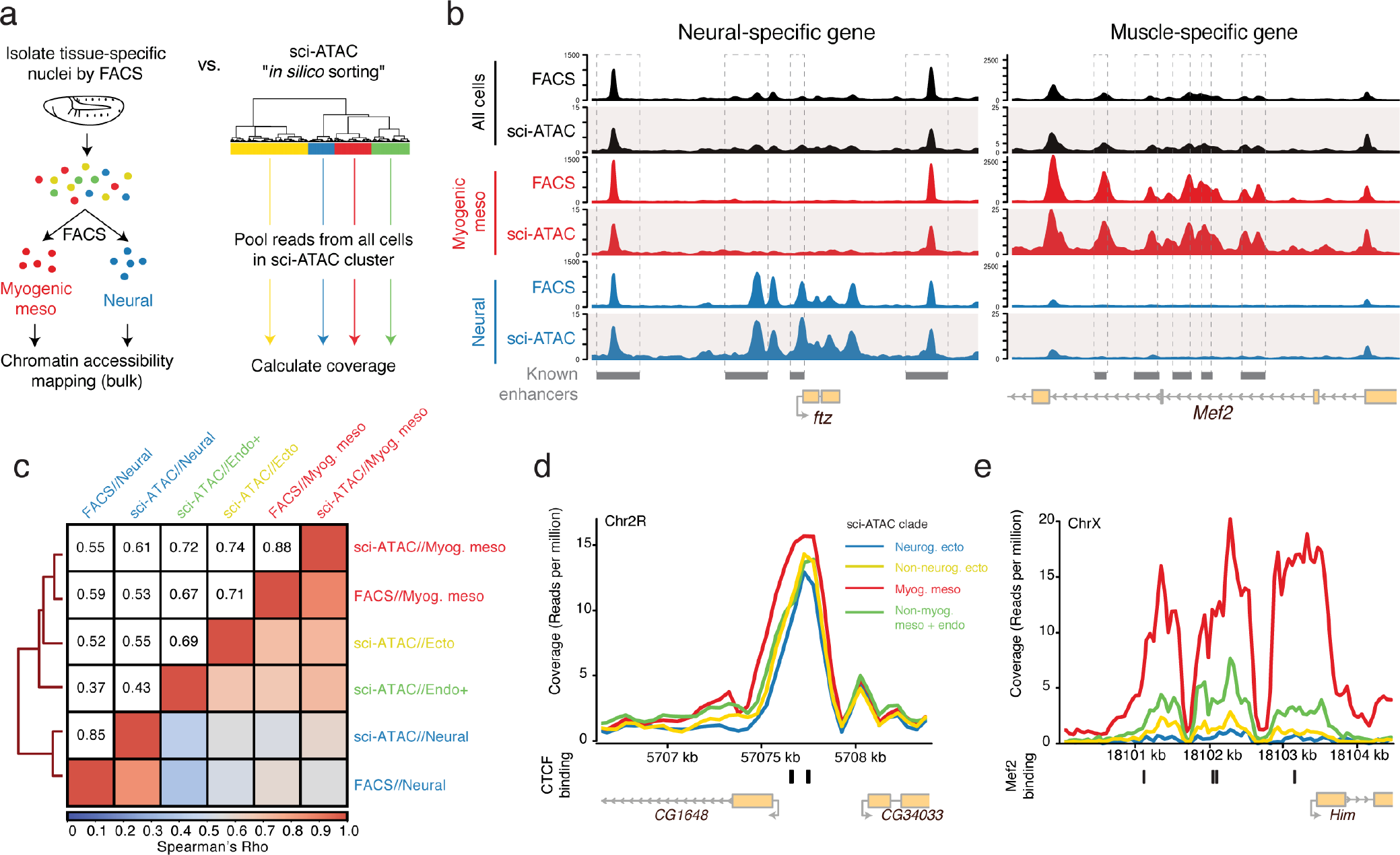
In *silico* sorting of single cell chromatin accessibility profiles. **a**, Contrasting FACS-DNase-seq and sci-ATAC-seq *“in silico* sorting.” Nuclei from embryonic mesoderm and neurons were isolated from 6-8 hr embryos using antibodies against tissue-specific regulatory proteins *mef2* (mesoderm) and *Elav* (neurons) followed by FACS and DNase. *In silico* sorts from sci-ATAC-seq were built by pooling reads from all cells within each LSI-defined clade. **b**, Library-size normalized coverage tracks from FACS-DNase (top in each color) and sci-ATAC-seq *in silico* sorts (bottom in each color) for whole embryo (black; whole-embryo DNase vs. all sci-ATAC-seq cells pooled), mesoderm (red; Mef2 DNase vs. sci-ATAC-seq clade 3), and neuronal (blue; Elav DNase vs. sci-ATAC-seq clade 1) at 6-8 hrs. Left, coverage around neural-specific transcription factor *ftz*. Right, coverage at intronic enhancers for mesoderm-specific transcription factor *mef2*. **c**, Cluster-gram and heatmap for all pairwise correlations (Spearman’s Rho) of normalized read counts at all distal peaks for the four sci-ATAC-seq *in silico* clades and the two FACS DNase datasets, all at 6-8 hrs. **d**, Normalized coverage plots for each sci-ATAC-seq clade over CTCF binding sites. All clades show highly similar accessibility. **e**, Normalized coverage plots for each sci-ATAC-seq clade over binding sites for the tissue-specific transcription factor *Mef2*. Read count density is considerably higher in the myogenic mesoderm (clade 3) and, to a much lesser extent in the clade comprised of non-myogenic mesoderm and endoderm (clade 4).

### Single cells cluster based on their germ layer of origin

To assess the identity of cells within each clade, we used two experimental sources of information (Fig. 1g): (1) the spatio-temporal activity of characterized developmental enhancers and (2) the tissue-specific expression of neighboring genes. The embryonic activity of approximately 4,000 *Drosophila* enhancers have been characterized *in vivo* with transgenics across all stages of *Drosophila* embryogenesis^4–6^. Similarly, the tissue-specific expression of ~60% of *Drosophila* genes has been characterized throughout embryogenesis by *in situ* hybridization^7,8^. We used these resources to determine if clade-specific promoter-distal elements (putative enhancers) were enriched for tissue-specific enhancer activity (Table S2) and whether clade-specific promoter-proximal elements (putative promoters) were enriched for genes with tissue-specific expression (Table S3).

For each clade, the enrichments of putative enhancers and promoters concurred, as did enrichments using all putative regulatory elements together (Table S4, Extended Data Fig. 3). The top enrichments show that the four major clades at the 6-8 hr and 10-12 hr time points largely correspond to the three major germ layers, with two subdivisions; ectoderm, which is split into neurogenic (clade 1) and non-neurogenic (clade 2) lineages, and mesoderm which is split into myogenic mesoderm (clade 3) and non-myogenic mesoderm (e.g. fat body and hemocytes) combined with endoderm (clade 4). The latter (clade 4) is interesting as it indicates that these tissues (non-myogenic mesoderm and endoderm) exhibit similar chromatin accessibility, a feature suggestive of a common developmental history. Although a common developmental origin is not known in *Drosophila*, this is highly reminiscent of the mesendoderm lineage in *C. elegans*^9^ and sea urchin^10^.

**Figure 3.**
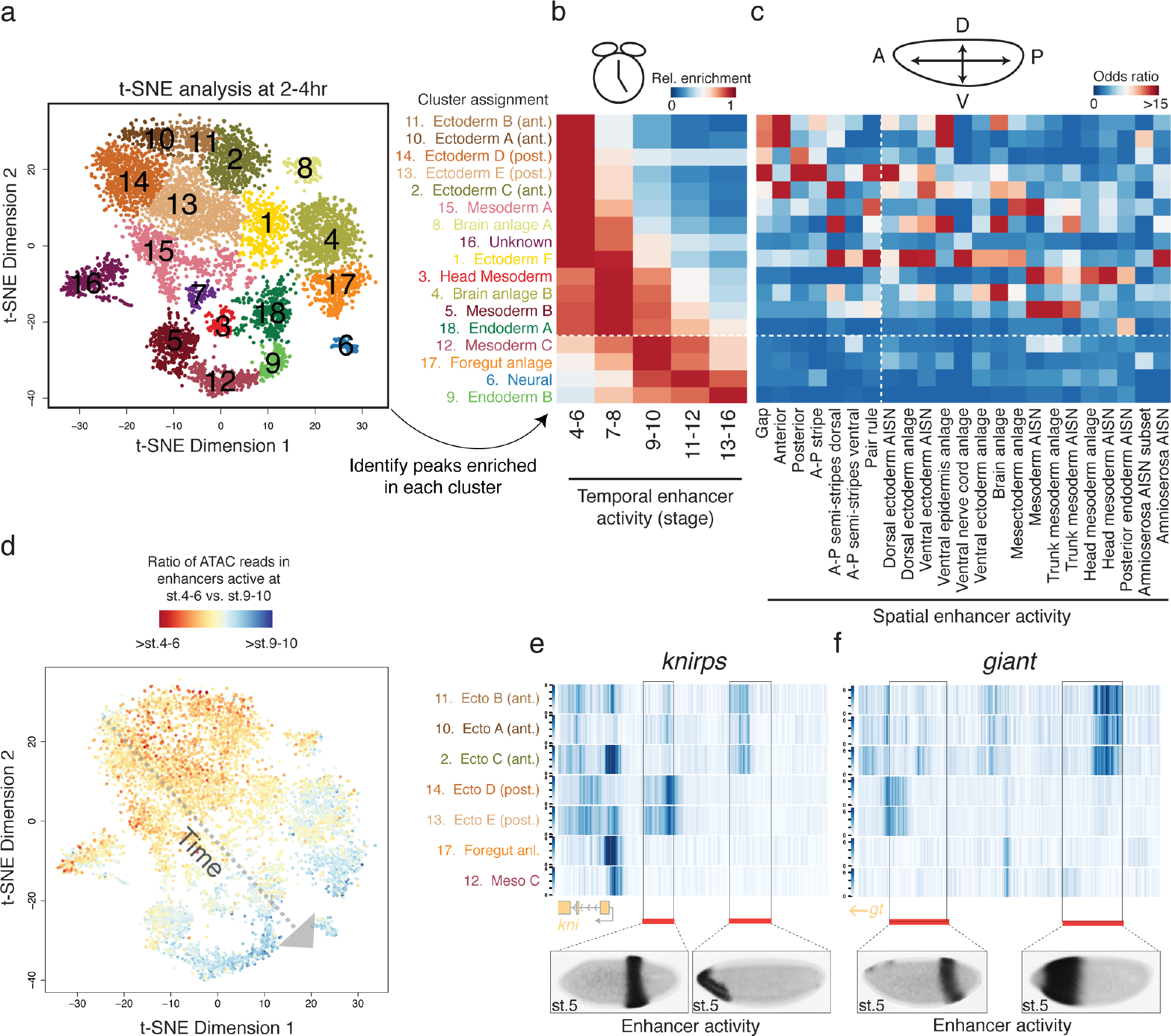
Temporal dynamics are a prominent driver of regulatory heterogeneity at 2-4 hrs of development. **a**, t-SNE clustering of sci-ATAC-seq data from the 2-4 hr time point. Cluster annotation was carried out by looking for overlaps between cluster-enriched peaks and enhancers with known tissue-specific activity (Methods). **b**, Relative enrichment of enhancers active at different developmental stages overlapping peaks enriched for read counts in the different 2-4 hr t-SNE clusters. Clusters below the white dashed line are likely derived from embryos outside of the 2-4 hr collection window, consequent to female ‘holding’ of older embryos. **c**, Enrichment (odds ratio) of enhancers active in different domains of the early embryo overlapping cluster-enriched peaks. Of note is a set of expression patterns (to the left of the dashed white line) corresponding to spatial patterns in pre-gastrulation embryos. **d**, Ratio of sci-ATAC-seq reads at enhancers active at stages 4-6 relative to reads at enhancers active at stages 9-10 for individual cells at 2-4 hrs. Reads in stage 4-6 enhancers are most enriched in cells appearing towards the upper left corner of the t-SNE plot. **e,f**, Heatmaps of library size normalized read counts in the vicinity of the gap genes *knirps* (e) and *giant* (f). In each case one characterized enhancer is known to drive anterior expression and another posterior expression in blastoderm embryos (stage5). *In situ* images of enhancer activity were obtained from^5^.

These clade assignments are further supported by enrichments for transcription factor (TF) binding motifs in clade-specific putative enhancers (Extended Data Fig. 4, Table S5). For example, at mid and late embryogenesis, motifs for the lineage specifying factors Krüppel (Kr), Tramtrack (ttk) and Runt (Run) were among the most enriched in the neurogenic ectoderm (clade 1), while Mef2 and CF2 motifs were most enriched in myogenic mesoderm (clade 3). GATA motifs were among the most highly enriched in the mesendoderm clade (clade 4) at 6-8 hrs and 10-12 hrs, reflecting the conserved role of GATA TFs in the specification of both non-myogenic mesoderm and endoderm^11,12^. In contrast, motifs for many TFs involved in antero-posterior and dorso-ventral spatial patterning of the early embryo were enriched at 2-4 hrs, including Hunchback (Hb), Bicoid (Bcd), Odd-paired (Odd) and Twist (Twi).

**Figure 4.**
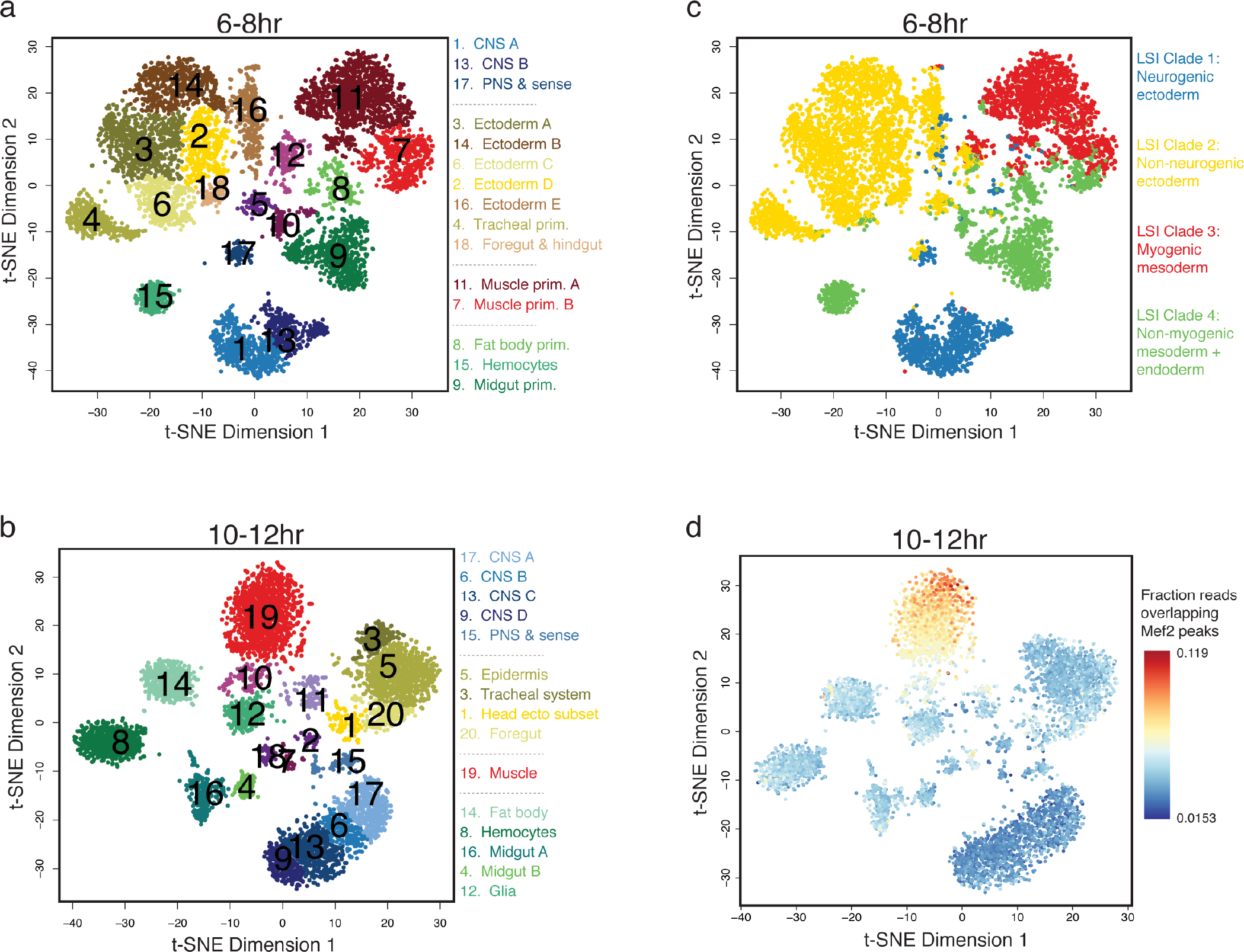
Single cells are readily assigned to tissues and cell types based on their chromatin accessibility. **a,b** t-SNE clustering of sci-ATAC-seq data from the 6-8 hr **(a)** and 10-12 hr **(b)** time point. Clusters were annotated based on overlaps between cluster-enriched peaks and enhancers with known tissue-specific activity (Methods). Three 6-8 hr (5, 10, 12) and five 10-12 hr (2, 7, 10, 11, 18) clusters are likely comprised of multi-cell ‘collisions’ based on library complexity and the distribution of reads mapping to the X chromosome (Methods; Extended Data Fig. 9). **c**, t-SNE clustered cells at 6-8 hrs colored by their original LSI clade assignment. The methods are largely consistent, with increased granularity in the t-SNE clustering. **d**, t-SNE clustered cells at 10-12 hrs, colored by the fraction of reads in each cell falling in Mef2 ChIP-seq peaks. There is a clear enrichment of reads at Mef2 binding sites in cluster 19 (muscle).

To experimentally evaluate how well our clade assignments correspond to germ layers and their subdivisions, we used FACS to isolate myogenic mesoderm and neuronal nuclei from 6-8 hr embryos using our previously published method^13^, obtaining ~98% purity for each. The FACS-isolated nuclei, as well as whole embryo samples, were subjected to DNase-seq in bulk, and the resulting maps of open chromatin compared to our *in silico* bulk (*i.e*. clade-defined) sci-ATAC-seq data at the same time point (Fig. 2a). The similarity between the resulting chromatin accessibility profiles for both methods for myogenic mesoderm (sci-ATAC-seq clade 3 vs. FACS-DNase) and neural ectoderm (sci-ATAC-seq clade 1 vs. FACS-DNase) are striking (Fig. 2b). For example, previously characterized neuronal enhancers near the *ftz* gene are accessible only in neurogenic ectoderm and not in myogenic mesoderm, with strong correlation between sci-ATAC-seq and FACS-DNase profiles (Fig. 2b, left panel). Similarly, enhancers at the *Mef2* locus are accessible in myogenic mesoderm but not neurogenic ectoderm, again with strong correlation between the two methods (Fig. 2b, right panel). This high agreement between both methods holds true globally, as seen by the high genome-wide correlations of open chromatin regions; >0.85 Spearman’s rho for both the myogenic mesoderm and neural ectoderm (Fig. 2c).

Tissue-specific enhancer activity is regulated by lineage-specific TFs such as Mef2 and Tinman for the myogenic mesoderm^14^, and Ttk and Kr for the neurogenic ectoderm, whose occupancy helps to define regulatory site usage^15^. Other factors such as CTCF and BEAF-32 play more widespread roles in genome organization^16^. To investigate the relationship between TF occupancy and chromatin accessibility within these clades, we compiled a set of probable binding sites for five well-studied mesoderm-specific transcription factors at 6-8hrs of development as well as sites for four TFs with more constitutive roles across tissues and developmental stages (*e.g*. CTCF and BEAF-32) (Extended Data Fig. 5). As expected, bound regions for ubiquitous factors such as CTCF exhibited similar levels of accessibility across all of the major clades (Fig. 2d, Extended Data Fig. 5a-d). In contrast, essential myogenic TFs (e.g. Tin, Mef2, Lmd) exhibited specificity for the myogenic mesodermal clade (Fig. 2e, Extended Data Fig. 5e-i). For example, three regions encompassing characterized mesoderm/muscle enhancers upstream of the muscle specific gene *Him* are bound by Mef2^17^ and have sci-ATAC-seq profiles that are strongly enriched within the myogenic mesoderm clade (Fig. 2e). These muscle-specific enhancers also exhibit notable accessibility in the non-myogenic mesoderm and endoderm clade (clade 4), albeit at lower levels than the myogenic mesoderm, likely reflecting a subset of non-myogenic mesoderm in which these Mef2 enhancers are also accessible.

**Figure 5.**
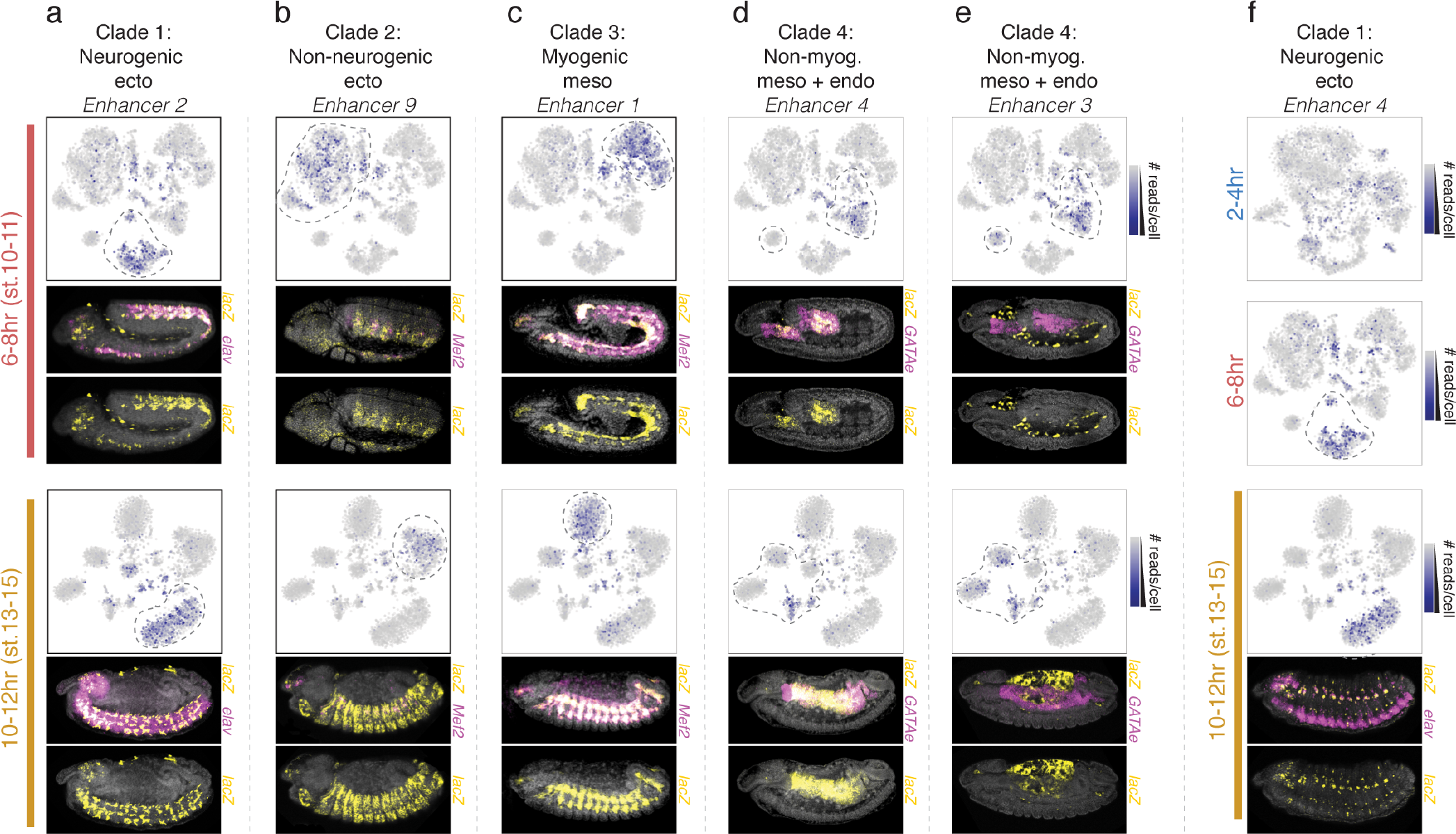
Prediction of tissue-specific enhancer activity using sci-ATAC-seq. **a-f**, Examples of candidate clade-specific enhancers tested in transgenic reporters. For each time point, upper panels shows the t-SNE plot for a specific time point with the intensity of blue representing the number of sci-ATAC-seq reads obtained from each tested element. Cell clusters bounded by dashed lines correspond to the predicted clade of activity. Lower panels show representative embryos for each time point with nuclei stained with DAPI (grey), and *in situ* hybridization for the reporter gene driven by the enhancer (yellow) and a tissue marker (magenta). The enhancer shown in (f) is first transcriptionally active at 10-12 hr (in the predicted tissue) but exhibits chromatin accessibility at earlier time points. All embryo images are lateral views, with anterior left and dorsal up. The activity of all tested enhancers is shown in Extended Data Fig. 12.

### Lineage-specific regulatory landscapes are established prior to gastrulation

Cells collected at the early time point (2-4 hrs) fall into five major clades (Fig. 1d), which are heterogeneous with respect to their chromatin accessibility. The clade identities were not immediately clear, in contrast to the later time points, potentially reflecting the plasticity of the early embryo where differentiated tissues have not yet formed. Moreover, these chromatin accessibility profiles are clearly distinct from later stages in embryogenesis, as seen when biclustering of cells from all three time points together (Extended Data Fig. 6). While cells from the 6-8 hr and 10-12 hr time points cluster together (stratified by germ layer), the cells from the 2-4 hr time point cluster separately.

Given this, we assessed whether these chromatin accessibility profiles corresponded to the earliest cell specification events and/or to anatomical patterning of the early embryo. Across its two-hour duration, 2-4 hr nuclei span embryos from the syncytial blastoderm, cellularization, gastrulation, and early germ-band extension stages of development (stages 5-8), with the majority of embryos (but not necessarily the majority of nuclei) being pre-gastrulation (stage 5). This is reflected in the general enrichment of the 2-4 hr clades for enhancers active in *either* early (stage 5) or late (stages 7-8) stages within this developmental window (Tables S2-S4), suggesting that temporal dynamics is a prominent driver of regulatory heterogeneity in this window. Developmental transitions occur very rapidly during these stages, with the pre-gastrula blastoderm (stage 5) lasting 40 minutes, and gastrulation onset (stage 6) and subsequent cell migration/division (stage 7) lasting only 10 minutes each. To capture finer granularity of cell clusters at this dynamic time window, we used ‘t-distributed stochastic neighbor embedding’ (‘t-SNE’), a dimensionality reduction method that has become popular in analyses of single cell RNA-seq data^18–20^. Specifically, we applied t-SNE to sci-ATAC-seq data using the master set of candidate regulatory elements (called from *in silico* aggregates of LSI-based clades), and then assigned cluster identities based on overlaps of enriched sites with enhancers with defined activity (Tables S6-S8). Because of confounding differences in sex chromosome copy number between male and female nuclei (Methods, Extended Data Fig. 7), we restricted the matrix to autosomal elements.

t-SNE applied to data from the 2-4 hr window yielded 18 cell clusters (Fig. 3a). Calculating the relative enrichment of active enhancers revealed marked differences in terms of each cluster’s developmental stage of maximal enrichment, allowing us to place them in an approximate temporal order (Fig. 3b). To further confirm that we are observing temporal heterogeneity, we calculated the ratio of ATAC reads per cell that fall into characterized enhancers known to be active at stages 4-6 vs. those known to be active at stages 9-10 (Fig. 3d, Extended Data Fig. 8a-d). Although t-SNE scaling should be interpreted very cautiously, we note a temporal axis from the top left to the bottom right of the plot (Fig. 3a,d). Eight clusters, mostly located towards the top left, are maximally enriched for stage 4-6 enhancers, as expected. In contrast, four clusters, appearing towards the bottom right, exhibit enrichment of enhancers active at stages, outside our collection window (Fig. 3b below the dashed line; clusters 12, 17, 6, and 9). These cells are likely contributed by a small proportion of older embryos (which have undergone more rounds of cell division) due to female ‘embryo holding’^21^. An additional four clusters exhibit maximal enrichment of stage 7-8 enhancers (clusters 3, 4, 5, 18), and appear between stage 4-6 and stage 9+ enriched clusters (Fig. 3a,d). Taken together, these observations suggest that temporal variation among single cells is a relatively large component of the total variance in chromatin accessibility at this early time point, and that dimensionality reduction using t-SNE can be used to assign relative developmental ages to cells based on their signatures of open chromatin. Interestingly, two of the developmentally early clusters were sex-biased (cluster 16: 85% male; cluster 8: 68% female; individual nuclei sexed based on ratio of autosomal to X chromosome reads; see Methods). We have been unable to determine the identity of the male-biased cluster, while elements specific to the female-biased cluster are enriched for brain anlage terms.

The 2-4 hr window is also when spatial patterning of the embryo along its antero-posterior (A/P) and dorso-ventral (D/V) axes is refined. We examined the relative enrichment of each 2-4 hr cluster for various categories of characterized developmental enhancers (Table S9) including those active early along one of the major embryonic axes, such as ‘posterior’ or ‘anterior’ (Fig. 3c, left of the vertical dashed line), or in the major territories of the early embryo or ‘anlagen’ that give rise to future embryonic structures or tissues (Fig. 3c, right of the dashed line)^5^. Most t-SNE clusters showed strong enrichment for multiple enhancer categories. Interestingly, nuclei with a developmentally early age (as defined in Fig. 3b,d; clusters 11, 10, 14, 13, 2, and 15) do not share a common chromatin accessibility landscape, but rather have varying enrichments for spatial enhancer terms. Chromatin accessibility within the loci of the two gap genes, *knirps* and *giant*, for example, varies in different cell clusters (Fig. 3e,f). The expression of *knirps* and *giant* are spatial patterned into broad strips along the A/P axis at this embryonic stage. Each of these genes has two characterized enhancers, one driving expression in a posterior stripe of the blastoderm and another in the anterior^5^. We identified several regions with differential accessibility between different 2-4 hr cell clusters: the anterior enhancers exhibit greater accessibility in the three clusters assigned to anterior presumptive ectoderm compared to the posterior presumptive ectoderm clusters, while the posterior enhancers exhibit higher accessibility in the posterior-assigned clusters (Fig. 3e,f). This implies that heterogeneity in chromatin accessibility among nuclei in different parts of the blastoderm embryo is established prior to gastrulation (Fig. 3a,c), and suggests a role for the early embryonic patterning systems in determining these differences. Consistent with this, many TFs known to be instrumental in patterning exhibit strong, differential enrichment across developmentally early t-SNE clusters (Extended Data Fig. 8e).

In addition to validating our t-SNE cluster assignments, these examples (Fig. 3e,f) illustrate how sci-ATAC-seq data can reveal spatially refined cells and their regulatory enhancers, without the need for FACS sorting. Interestingly, classic lineage-tracing and transplantation experiments showed that the broad fate, and to some extent, the developmental potential of cells is largely determined at the cellular blastoderm stage, leading to the idea of a blastoderm fate map^22^. Our data suggest that these early cell specification events are underpinned by regional differences in chromatin accessibility.

### A fine-grained map of single cell tissue identities

We next applied t-SNE as described above to the later two time points, with the goal of resolving individual cell types and tissues during their lineage commitment (6-8 hrs) and differentiation (10-12 hrs). The vast majority of cell clusters could be readily assigned to specific tissues or cell types at these later time points (Fig. 4a,b) based on their enrichments for tissue specific enhancers (as described above; Tables S6-S8), while a number of small clusters (Fig. 4a,b; purple clusters) were discarded as they are likely ‘collisions’ resulting from our combinatorial indexing methodology (Methods; Extended Data Fig. 9)^1^. These tissue or cell type assignments are broadly consistent with the germ layer assignments of LSI-based clades (Fig. 1, Extended Data Fig. 10), but with much finer granularity (Fig. 4a,b). For example, the non-myogenic mesoderm and endoderm (clade 4 in Fig. 1, Fig. 4c) is resolved into three separate clusters at 6-8 hrs (and five at 10-12 hrs) comprising the fat body (cluster 8) and hemocytes (cluster 15) from the non-myogenic mesoderm, and midgut (cluster 9) from the endoderm (Fig. 4a). These tissue assignments also match the embryonic occupancy of transcription factors with known roles in the tissues identity or progression (Tables S6-S8). For example, cells belonging to cluster 19 (muscle) at 10-12 hrs show a higher proportion of their sci-ATAC-seq reads mapping to embryonic chromatin immunoprecipitation (ChIP) peaks for the key myogenic transcription factor Mef2 than cells in other clusters (Fig. 4d).

To further explore the biological properties of each cluster, we combined reads from cells within each t-SNE cluster to generate tissue or cell type-specific “*in silico* bulk” tracks of chromatin accessibility at each time point. Contrasts among these tracks revealed a wealth of differences between clusters that could not have been distinguished in bulk data without substantial efforts to FACS-isolate cells using transgenic markers or antibodies to lineage specific nuclear factors. For example, the neural clusters at 10-12 hrs show extensive differences in accessibility surrounding genes involved in neural specification, highlighting both spatial, and potentially temporal, differences in the regulatory landscape among neural precursors over the course of differentiation (Extended Data Fig. 11).

### sci-ATAC-seq accurately predicts the activity of tissue-specific enhancers

A major advantage of chromatin accessibility profiling over transcriptome profiling is its potential to identify the regulatory elements that drive single cell trajectories during development. We therefore asked whether elements exhibiting tissue-specific chromatin accessibility during embryogenesis corresponded to bona fide tissue-specific enhancers. We selected 31 putative enhancers; promoter-distal elements exhibiting strong clade-specific accessibility at 6-8 hrs and/or 10-12 hrs, that do not overlap previously tested regions in transgenic enhancer assays (see Methods; Table S10). Of note, other than clade-specific accessibility, no other criteria were applied to bias the selection towards different kinds of activity (*e.g*. enhancers vs. insulators, for example). Six of the candidate regions exhibited specific accessibility in neurogenic ectoderm, ten in non-neurogenic ectoderm, eight in myogenic mesoderm, and seven in non-myogenic mesoderm plus endoderm. Each putative regulatory element was cloned in front of a minimal promoter driving a *lacZ* reporter gene and stably integrated into a common location in the *Drosophila* genome using the Phi-integrase system to minimize positional effects. Transgenic embryos were collected across all stages of embryogenesis and enhancer activity was assessed by *in situ* hybridization against the *lacZ* reporter together with a specific tissue marker gene.

Surprisingly, given the simple strategy used to select elements, 93% (29/31) of candidate regions function as developmental enhancers *in vivo* (Fig. 5; Table 1; Extended Data Fig. 12; Table S10). Remarkably, all 29 active enhancers showed activity in the tissue we predicted based on our sci-ATAC-seq data; 26 were exclusive to the predicted tissue, while three elements drove activity in small subsets of cells from other clades in addition to the predicted tissue (Table 1; Table S10). For example, elements with accessibility specific to either the ectodermal, muscle or mesendodermal clades show enhancer activity exclusively in the developing epidermis (Fig. 5b), muscle (Fig. 5c), or midgut (Fig. 5d), respectively, perfectly matching their predicted activity. In contrast, an element with predicted activity in the nervous system (clade 1) is active in both the central nervous system and a small subset of cells in the amnioserosa (clade 2) (Fig. 5a).

**Table 1.** Overview of enhancer activity assay results for a set of predicted clade-specific regulatory elements.

|  | Neurogenic<br>ectoderm | Non-neurogenic<br>ectoderm | Myogenic<br>mesoderm | Non-myogenic<br>mesoderm and<br>endoderm | Total |
| --- | --- | --- | --- | --- | --- |
| **Active as**<br>**enhancer** | 4/6 (67%) | 10/10 (100%) | 8/8 (100%) | 7/7 (100%) | 29/31 (94%) |
| **Correct tissue**<br>**(inclusive)** | 4/4 (100%) | 10/10 (100%) | 8/8 (100%) | 7/7 (100%) | 29/29 (100%) |
| **Correct tissue**<br>**(exclusive)** | 3/4 (75%) | 10/10 (100%) | 7/8 (88%) | 6/7 (86%) | 26/29 (90%) |

Importantly, tested elements enriched in the non-myogenic mesoderm and endoderm clade often act as enhancers in either hemocytes or gut endoderm, showing distinctions in the regulatory landscapes of these germ layers. Enhancer 4, for example, has enriched open chromatin in cells of the developing midgut at both time points, in concordance with the enhancer’s specific activity in the developing anterior and posterior midgut during these stages of embryogenesis (Fig. 5d). Notably, three out of the seven tested enhancers drove specific activity in yolk nuclei (Fig. 5e, Extended Data Fig. 12d). As the yolk is mainly extra-embryonic, this was unexpected and suggests a potential regulatory link between the yolk and mesendodermal tissues.

Interestingly, the patterns (both temporal and spatial) of accessibility and observed activity are not always perfectly synchronous. In some cases the enhancer’s accessibility is broad, spanning multiple t-SNE clusters, yet the enhancer is only active in a subset of predicted tissues. The non-myogenic mesoderm and endoderm enhancer 3, for example, has broad accessibility in midgut, fat-body and hemocytes, but its activity is restricted to yolk nuclei (Fig. 5e). This may reflect alternative TFs binding to the same enhancer in different tissues, or possibly activators binding in yolk cells and repressors in the other tissues. We also noted that in some instances the enhancer was accessible much earlier (and again in a broader range of cell types) than when the enhancer was first detected to be active in the transgenic embryos, suggesting that these regulatory elements may be “primed” before they actually influence target gene expression. For example, the neurogenic ectoderm enhancer 4 only initiates activity at stage 13 specifically in the peripheral nervous system and then weakly in the brain, yet the enhancer is accessible in neuronal cell clusters, and also in some non-neuronal clusters, already at 2-4 hrs (Fig. 5f).

In summary, these results demonstrate the power of sci-ATAC-seq not only to elucidate the dynamics of chromatin accessibility between cell types and across time, but also for the large-scale prediction of *in vivo* enhancer activity. Altogether, this study identified 30,499 putative distal regulatory elements exhibiting clade-specific accessibility (Table S1), 18% of which overlap previously characterized developmental enhancers, and 51% of which overlap putative enhancers as defined by bulk embryonic DHS data^3^. However, in contrast with the *in silico* sorting of single cell ATAC-seq data, bulk DHS from whole embryos fails to pinpoint the tissue and cell-type specificities of regulatory elements. This genome-wide database of putative regulatory elements and their predicted cell type and temporal specificities, represents a powerful resource for future investigations of *Drosophila* embryogenesis.

## DISCUSSION

The past four decades have seen vast advances in our understanding of the molecular pathways leading to spatial patterning and tissue differentiation in the developing embryo. These insights are largely based on detailed investigations of: (1) gene expression patterns via *in situ* hybridization and immunostaining, and (2) gene function via genetic loss- and gain-of-function coupled to imaging-based phenotyping. Although by their very nature these methods examine single embryos and single cells, they do not readily ascertain global views of the regulatory landscape as it unfolds during development.

Early efforts to examine the molecular underpinnings of embryonic cell lineages at a genome-wide scale have relied on marker-based cell sorting via FACS or microdissection. However, such studies are necessarily based on prior knowledge and can only examine a particular cell type after a pre-defined lineage marker is expressed. In contrast, the “shotgun” analysis of single cells represents an unbiased (*i.e*. marker-free) strategy for characterizing the genomic landscape of biological systems, including the molecular trajectories that underpin the differentiation of cell types. With the recent advent of highly scalable methods, the opportunity is ripe to obtain global, single cell views of well-studied systems such as *Drosophila* embryogenesis.

This study represents the first application of single cell chromatin accessibility profiling to the *in vivo* development of a multicellular organism. Chromatin accessibility is a powerful marker of regulatory DNA. The enhanced version of sci-ATAC-seq that we present here obtains tens of thousands of reads per cell, and can readily process tens of thousands of cells per experiment. In this study, we present chromatin accessibility data for ~23,000 single cells spanning three major stages of *Drosophila* embryogenesis. These data provide a global view of the changes in chromatin accessibility as an embryo transitions from a pre-gastrula stage through to cell lineage commitment and subsequent terminal tissue differentiation. Our results reveal groups of cells with shared profiles of open chromatin that reflect their tissue/cell type in older embryos, and their anatomical position in younger pre-gastrula embryos. Rather than observing a shared chromatin landscape in blastoderm embryos, our data reveals clusters of cells with distinct patterns of open chromatin that likely reflect the segmentation of the early embryo along its A/P and D/V axis.

The identification of tissue-specific enhancers has been a major challenge for the field of genomics. We show here that elements exhibiting tissue-specific chromatin accessibility during *Drosophila* embryogenesis, as identified through the lens of shotgun sampling of single cells followed by *in silico* sorting, are overwhelmingly likely to function as developmental enhancers when validated *in vivo* in transgenic reporter assays. We anticipate that the large-scale profiling of chromatin accessibility in early development in mouse and models of human development^23^ may facilitate the global discovery of developmental enhancers involved in mammalian embryogenesis.

Looking forward, this study motivates at least three challenging, but essential, lines of further study in *Drosophila*. First, single cell chromatin accessibility profiling through the entirety of fly development, with the goal of defining a fully continuous view of the regulatory landscape as it unfolds. Second, applying sci-ATAC-seq as a ‘phenotype’ in place of imaging, *i.e*. to take advantage of the wealth of characterized mutants available in *Drosophila* to dissect the requirement of different TFs and chromatin regulators in the regulatory decisions of single cells during embryogenesis. Finally, concurrently measuring chromatin accessibility, transcription, and cell lineage history^24,25^ in single cells during embryogenesis, which together with the above will advance us towards a truly comprehensive understanding of how the genome encodes metazoan development.

## METHODS SUMMARY

We optimized sci-ATAC-seq to obtain roughly an order of magnitude more reads per cell than previously reported^1^, and to facilitate use with formaldehyde fixed nuclei. For three time points in embryogenesis (2-4 hrs, 6-8 hrs, and 10-12 hrs), we generated pooled sci-ATAC-seq libraries (information from each individual cell is distinguished by a unique barcode) that were sequenced on a NextSeq sequencer (Illumina). All mapped reads were filtered to remove non-unique reads, ambiguous barcodes and duplicates. Data were first evaluated by latent semantic indexing (LSI). To do so, 2 kb genomic bins were scored for accessibility per cell per time-point, removing cells with the lowest 10% of accessible sites. This large binary matrix was normalized and re-scaled using term frequency-inverse document frequency (“TF-IDF”) transformation followed by singular value decomposition to generate a lower dimensional representation of the data by only considering the 2nd through 6th dimensions. These LSI scores of accessibilities were used to cluster cells and windows based on cosine distances. To call open chromatin peaks within each cluster, uniquely mapped reads per cell were aggregated across all cells in a cluster and peaks called using MACS2^26^. Cell types were identified through enrichments of characterized tissue-specific enhancers, using a custom database of ~8,000 enhancers tested in transgenic embryos, in addition to the near-by gene’s expression (for promoter proximal elements). To refine our cluster definitions, we used t-SNE and density peak clustering. After transformation of the data with TF-IDF, we generated a lower dimensional representation of the data by including the first 50 dimensions of the singular value decomposition, which was used as input for the Rtsne package^18,27,28^. Clusters were defined by the density peak clustering algorithm in Monocle 2^29^.

All raw data will be submitted and released through ArrayExpress and GEO. BigWig files for coverage within each clade, regions of accessibility (peak calls), and a master list of all potential regulatory elements (Table S1) will be made available on the Furlong lab web page http://furlonglab.embl.de/data. To make the data easily accessible we have generated a searchable html where the user can select a tSNE cluster or genomic locus of interest and visualize the data throughout the genome that will also be made available.

## ACKNOWLEDGEMENTS

This work was technically supported by the EMBL Advanced Light Microscopy Facility. We thank D. Prunkard, and L. Gitari in the UW Pathology Flow Cytometry Core Facility for their exceptional assistance in flow sorting. We thank all members of the Furlong and Shendure laboratories for discussions and comments. This work was financially supported by EMBL and BMBF (TransDiag 2) funds to EEF and NIH (DP1HG007811 and R01HG006283) and the Paul G. Allen Family Foundation funds to JS. DAC was supported in part by T32HL007828 from the National Heart, Lung, and Blood Institute. JS is an Investigator of the Howard Hughes Medical Institute.

---

**Extended Data Figure 1.**
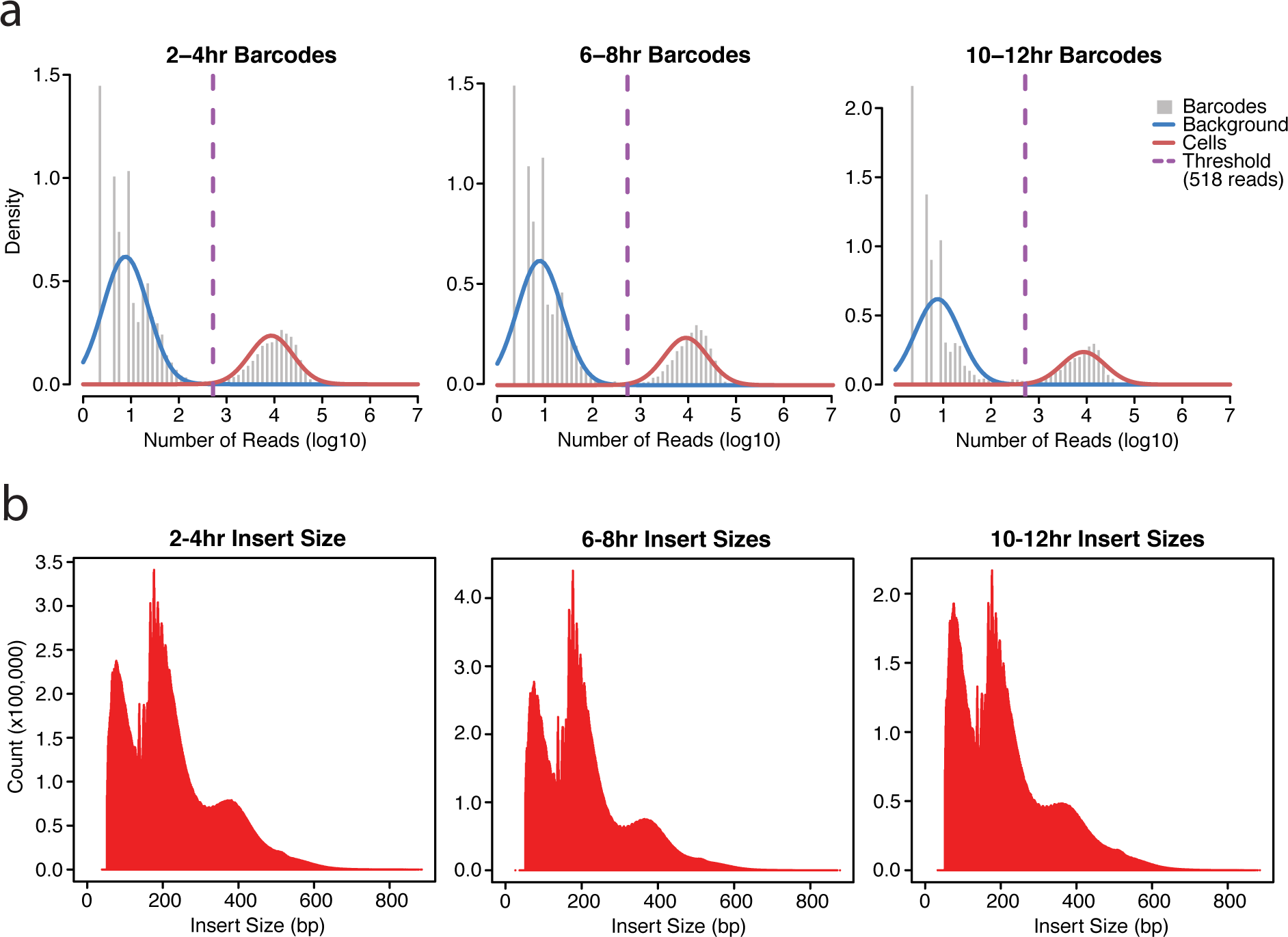
Distribution of read counts and fragment size per cell. **a**, Log10 counts of sci-ATAC-seq reads per barcode at each time point show a bimodal distribution. The bimodal distribution was used to set a threshold of 518 reads to identify barcodes corresponding to valid cells vs. background. **b**, Fragment size distribution at each of time point is consistent with the nucleosomal banding pattern observed in standard (bulk) ATAC-seq experiments.

**Extended Data Figure 2.**
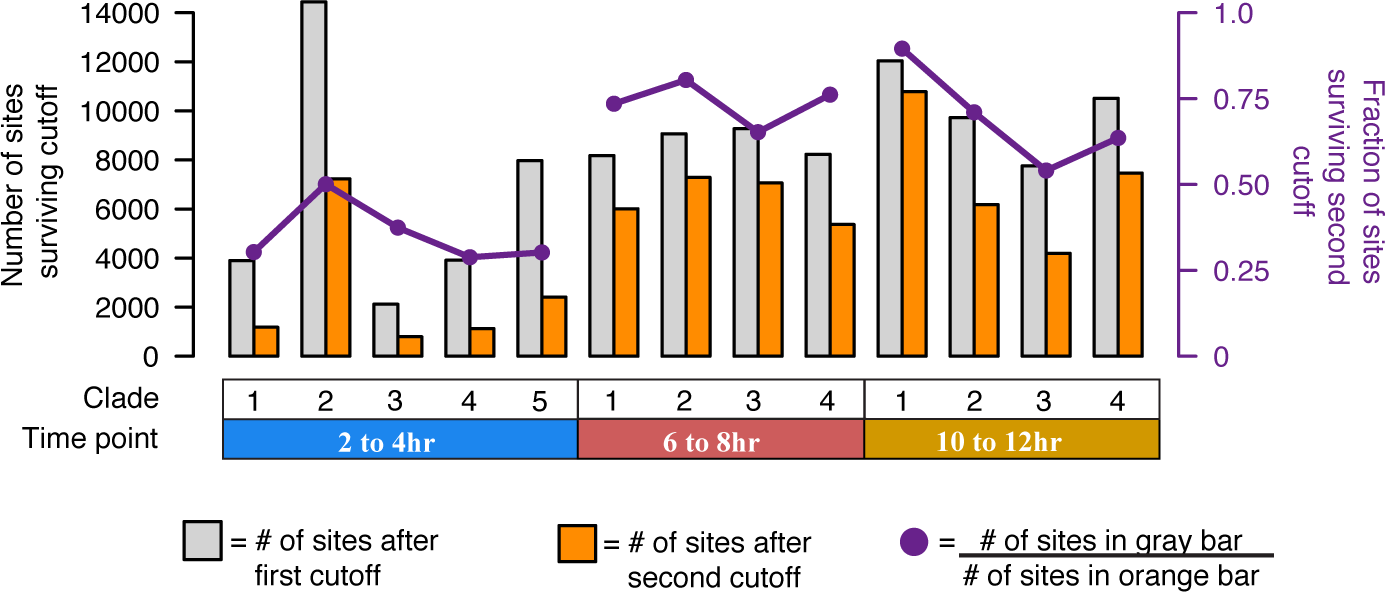
Number of clade-specific sites identified at each time point. Barplot of the number of sites identified as significantly open in each clade (1% FDR; gray bar) and the number of sites specific to that clade (Methods; orange bar). Overlaid on the barplot (purple points) is the fraction of sites passing the first threshold that also pass the second threshold (count of orange bar / count of gray bar).

**Extended Data Figure 3.**
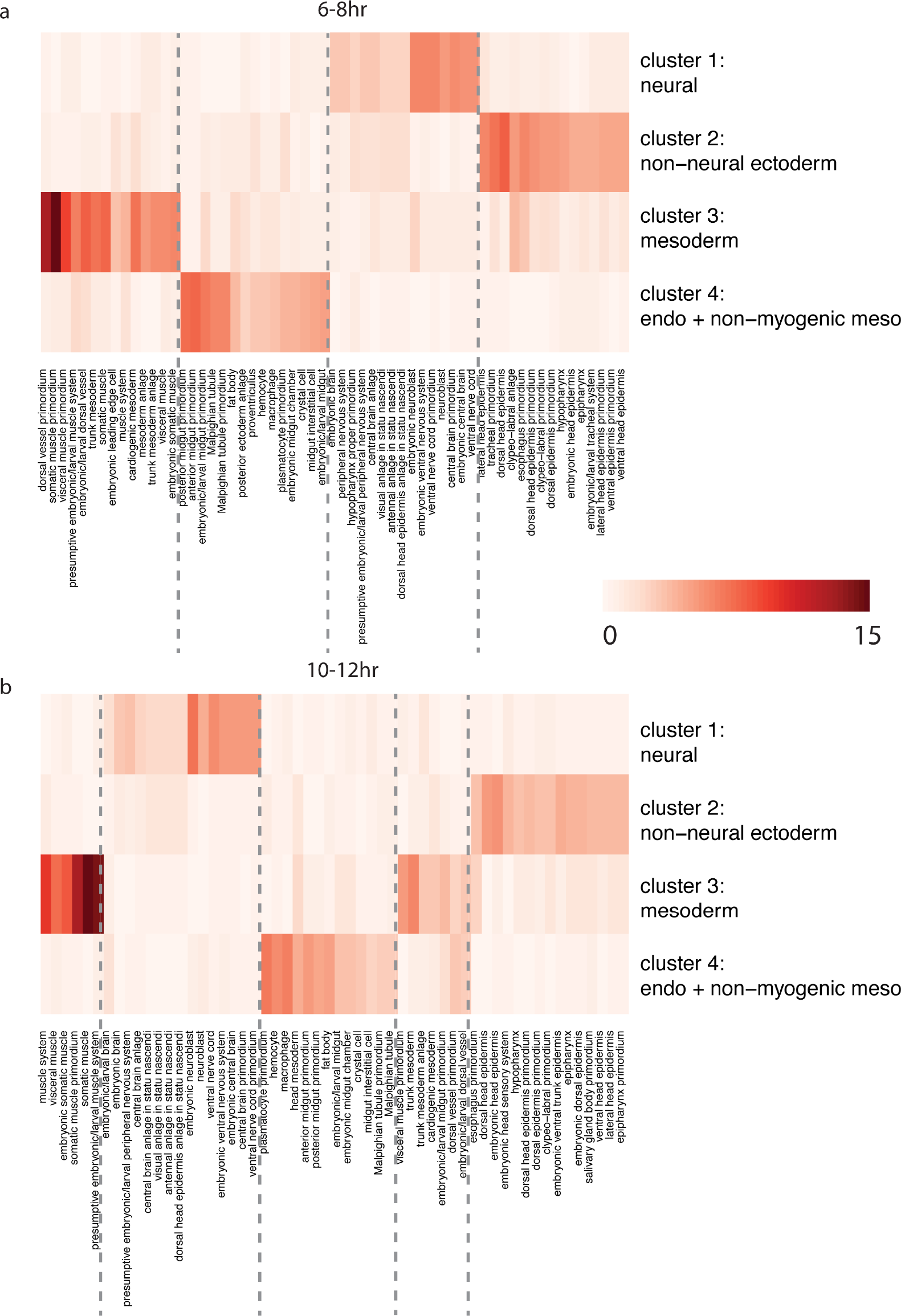
Enhancer enrichments for LSI clades at 6-8 hr sand 10-12 hrs. Enrichment for tissue-of-expression information for characterized distal enhancers overlapping clade-specific peaks at 6-8 hrs and 10-12 hrs of development. Shown are all categories in the top 10 enrichment of any clade (enrichment scores capped at 15 for display) containing at least 20 known enhancer overlaps. **a**, Enrichments of distal enhancer expression terms at 6-8 hrs after egg laying. **b**, Enrichments of distal enhancer expression terms at 10-12 hrs after egg laying.

**Extended Data Figure 4.**
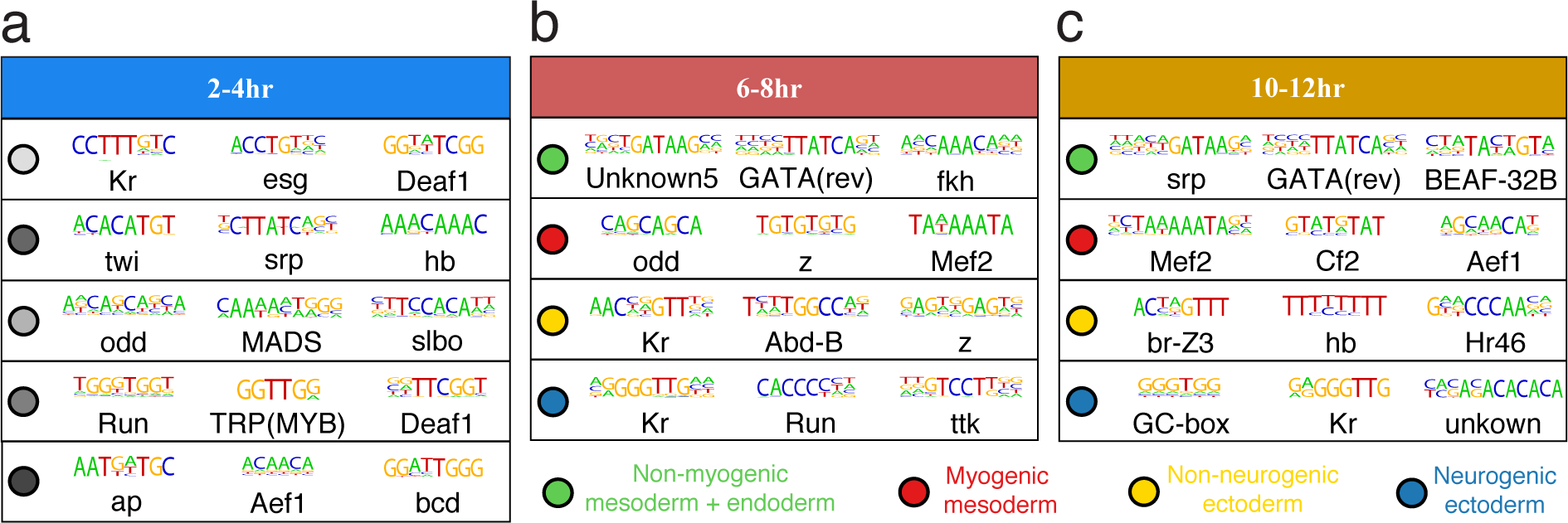
Motifs enriched in peaks LSI clade-specific peaks. SeqGL was run on LSI clade-specific distal peaks at each time point to identify enriched sequence motifs. The top 3 most enriched motifs for each clade are displayed. Colored circles indicate which clade is represented by each line. For the later time points (6-8 hrs and 10-12 hrs), green is endoderm, red is myogenic mesoderm, yellow is non-neurogenic ectoderm, and blue is neurogenic ectoderm. The results show an enrichment of motifs for factors associated with early development at 2-4 hrs with more tissue-specific factor motifs (e.g. mesodermal factor Mef2 or neural regulator Tramtrack (ttk)) within germ-layer annotated clades at later stages of development.

**Extended Data Figure 5.**
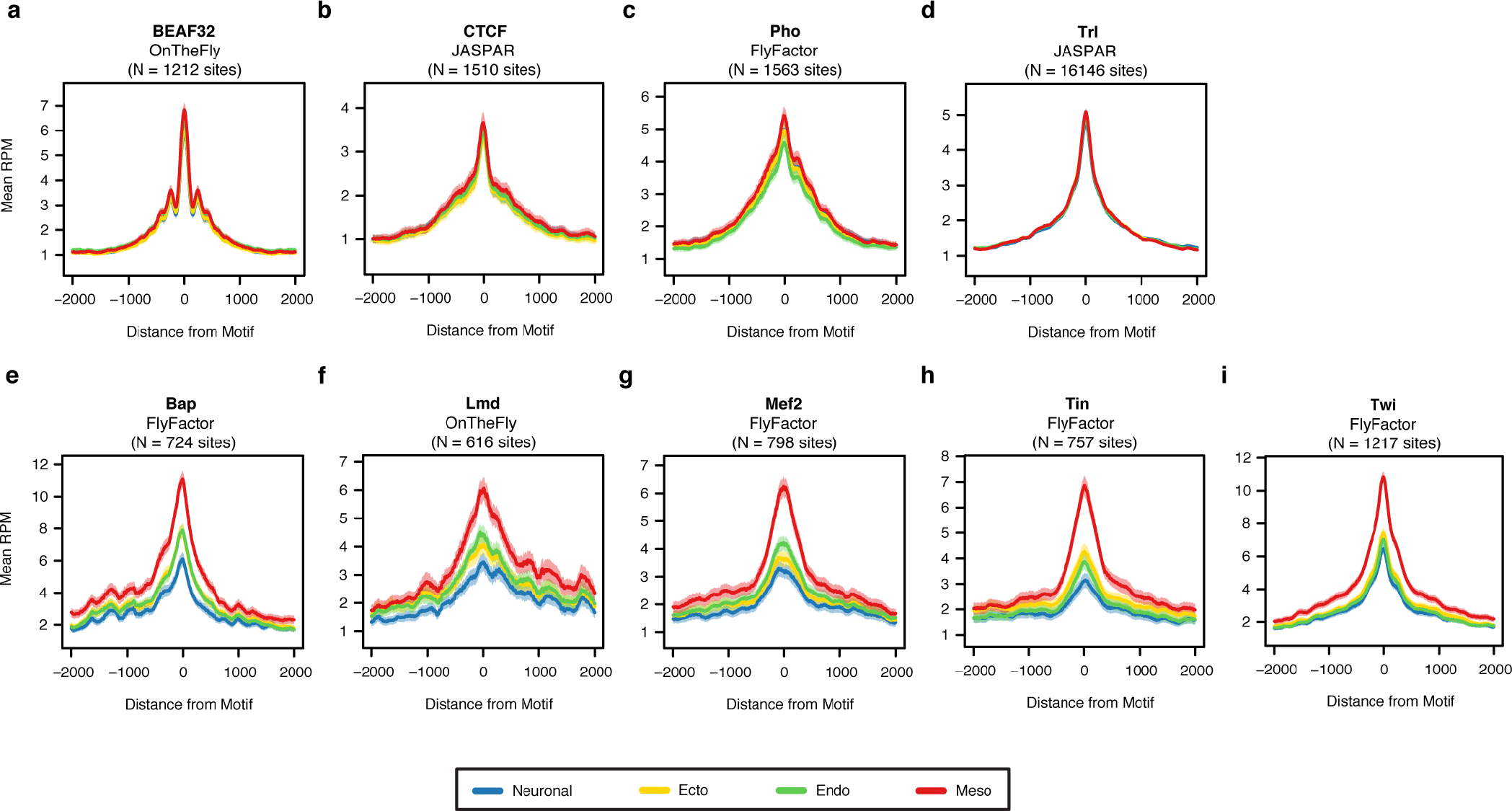
Chromatin accessibility at mesoderm-specific versus ubiquitous TF binding sites for each LSI clade. Using ChIP occupancy data (peaks) and transcription factor binding motifs compiled previously^32^, we scanned for all TF motif instances under ChIP peaks from datasets spanning 6-8 hrs of development using FIMO. Aggregate read counts in 4kb windows around each identified motif instance is shown for each of the four LSI clades at 6-8 hrs. Green is endoderm, red is myogenic mesoderm, yellow is non-neurogenic ectoderm, and blue is neurogenic ectoderm. **a-d**, Aggregate plots for four ubiquitous transcription factors (BEAF32, CTCF, Pho, and Trl) at 6-8 hrs. **e-i**, Aggregate plots for mesodermal transcription factors (Bap, Lmd, Mef2, Tin, Twi) at 6-8 hrs.

**Extended Data Figure 6.**
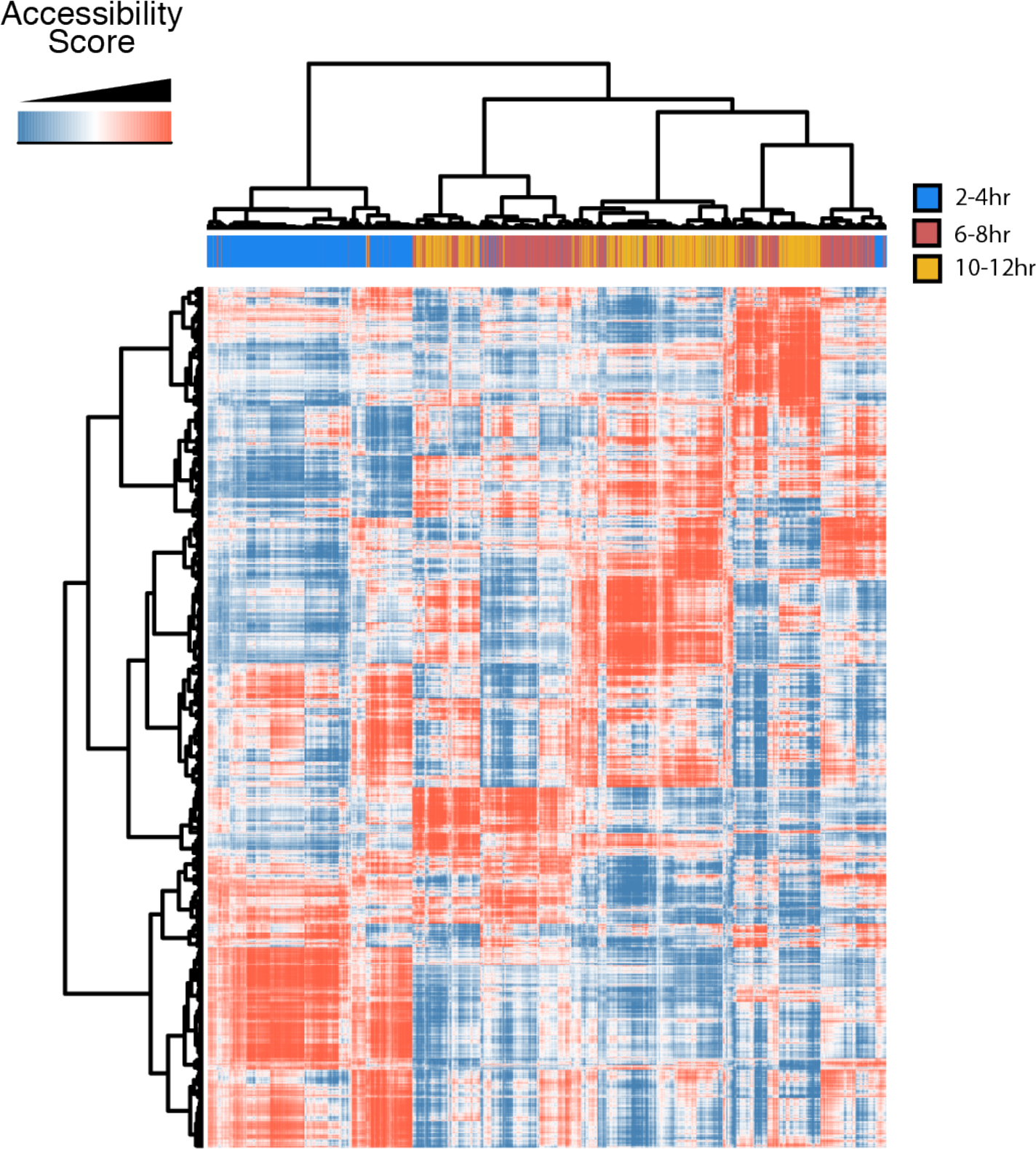
Clustered LSI data for all three time points processed together. In addition to processing data from each time point independently, data from all cells can be analyzed together (with the caveat that time point and batch are now confounded). Here, we show binarized, LSI-transformed, and clustered count data for 2 kb windows across the genome for cells from all three time points (blue = 2-4 hrs, red = 6-8 hrs, orange = 10-12 hrs) processed together. The predominant pattern is one in which pre-gastrulation cells (2-4 hrs) cluster together while post-gastrulation cells (6-8 hrs and 10-12 hrs) cluster together roughly by germ layer-of-origin with four main clades present in both the 6-8 hr and 10-12 hr samples.

**Extended Data Figure 7.**
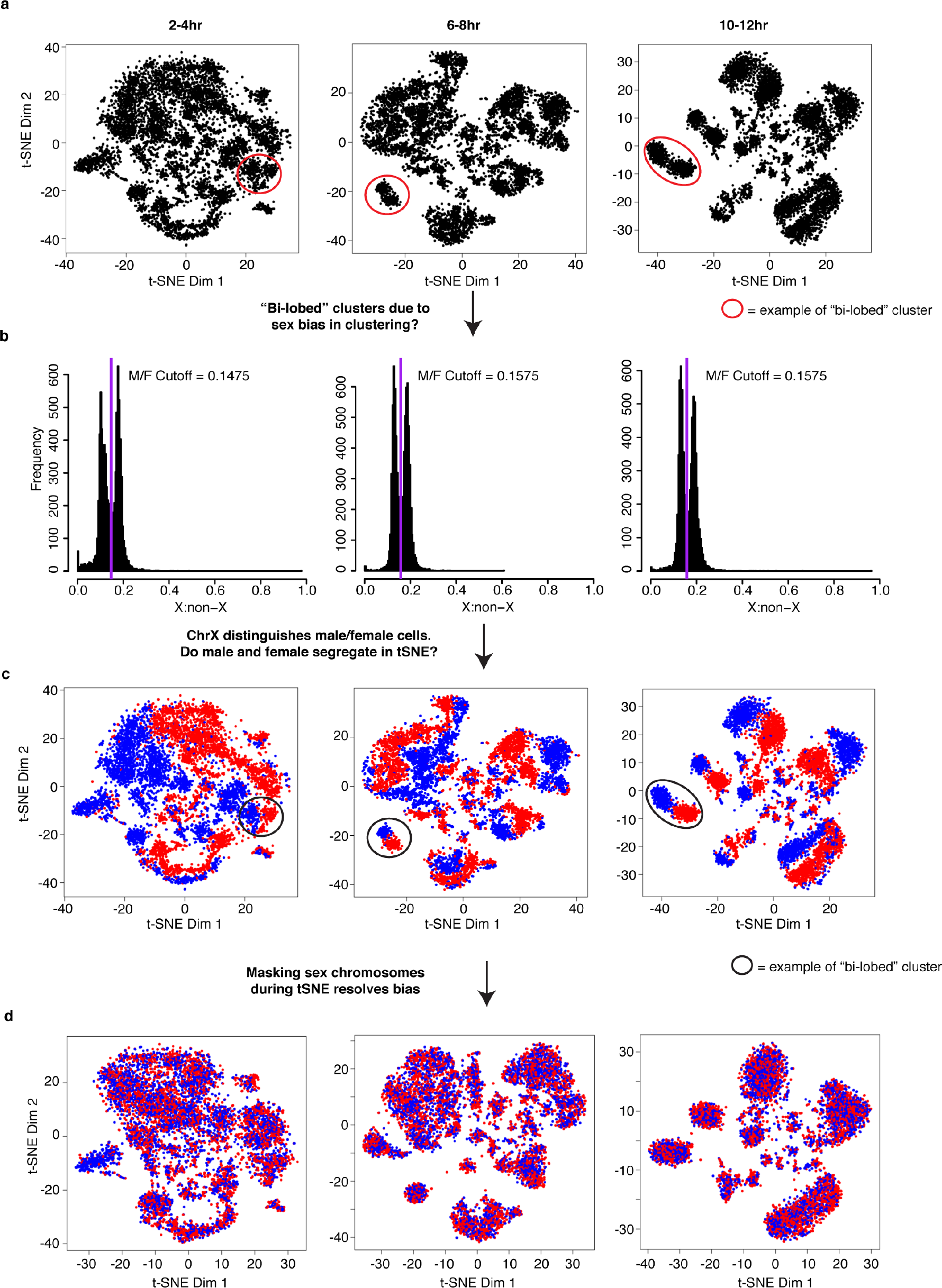
Sex of individual cells identified by ratio of X:autosomal reads. Embryos at all stages consist of a mixture of male and female embryos (males: XY, females: XX). **a**, t-SNE plots of three time points from our initial analysis. Many clusters exhibited a “bi-lobed” structure, where each individual cluster was made up of two “mirrored” lobes (red circles identify one example of bi-lobed clusters from each time point). This was most apparent at the 10-12 hr time point. **b**, Histogram of the ratio of chromosome X to autosomal reads in individual cells. To explore whether this “bi-lobed” structure was a function of sex biases in clustering, we attempted to sex individual cells. The ratio of X:autosomal reads shows a bimodal distribution as expected in a system with heterogametic (XY) males and no evidence for imprinting. The purple line marks the local minimum between the two peaks of the histograms. **c**, Initial t-SNE clusters colored by sex assignment. Coloring individual cells by their sex reveals that the “bi-lobed” architecture is largely driven by sex biases in clustering. **d**, After removing X chromosomal reads, data was re-clustered and individual cells recolored by the ratio of X:autosomal reads. The resulting clusters showed an approximately equal number of male and female cells except for clusters 8 and 16 at the 2-4 hr time point.

**Extended Data Figure 8.**
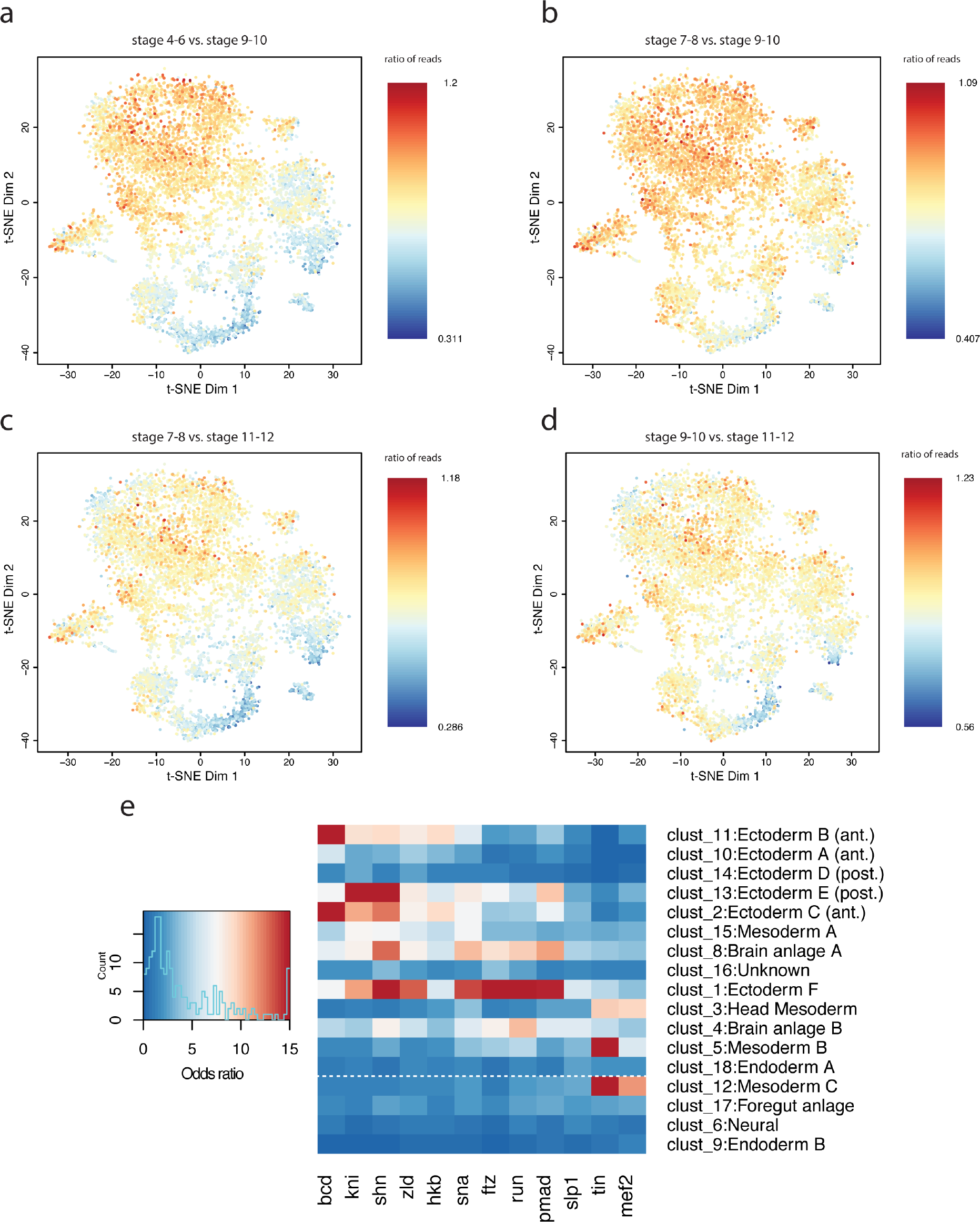
t-SNE reveals that temporal and spatial heterogeneity of cells at the 2-4 hr time point is reflected in chromatin accessibility. **a-d**, Ratio of total read counts per cell for enhancers active at different stages of development. Read counts within temporally characterized enhancers provide insight into the specific stage of development from which a cell is derived. Plotted here are ratios of counts in earlier vs. later active enhancers showing a rough temporal progression of cells from upper left to bottom right with cells from the earliest embryonic stages (stages 4-6) clustering in the top left. **a**, Ratio of reads in enhancers active at stages 4-6 vs. those active at stages 9-10 (as in Fig. 3d). Cells are enriched for reads active in stages 4-6 in clusters appearing towards the top left. **b**, Ratio of reads in enhancers active at stage 7-8 vs. stage 9-10 showing a shift in the enrichment towards the bottom right. **c**, Ratio of reads in enhancers active at stages 7-8 vs. stage 11-12 (contrast with (a)). **d**, Ratio of reads in enhancers at stages 9-10 vs. stage 11-12 (contrast with (a)). **e**, Enrichments (odds ratio) of transcription factor ChIP peaks overlapping t-SNE cluster-enriched peaks at 2-4 hrs. Data were taken from Table S9 and reflect only ChIP data from appropriately staged embryos. Plotted here are only those transcription factors (columns) whose maximum enrichment across the t-SNE clusters were at least 5. Temporally earlier clusters show a clear enrichment of peaks for TFs conferring spatial positioning information in the early embryo. Cells assigned to later stages (e.g. clusters 17, 6, and 9) show relatively low enrichment for early embryo ChIP peaks (an exception is cluster 12 containing cells from mesodermal tissues in slightly older embryos, which show a continued enrichment for mesodermal regulatory factors Tin and Mef2).

**Extended Data Figure 9.**
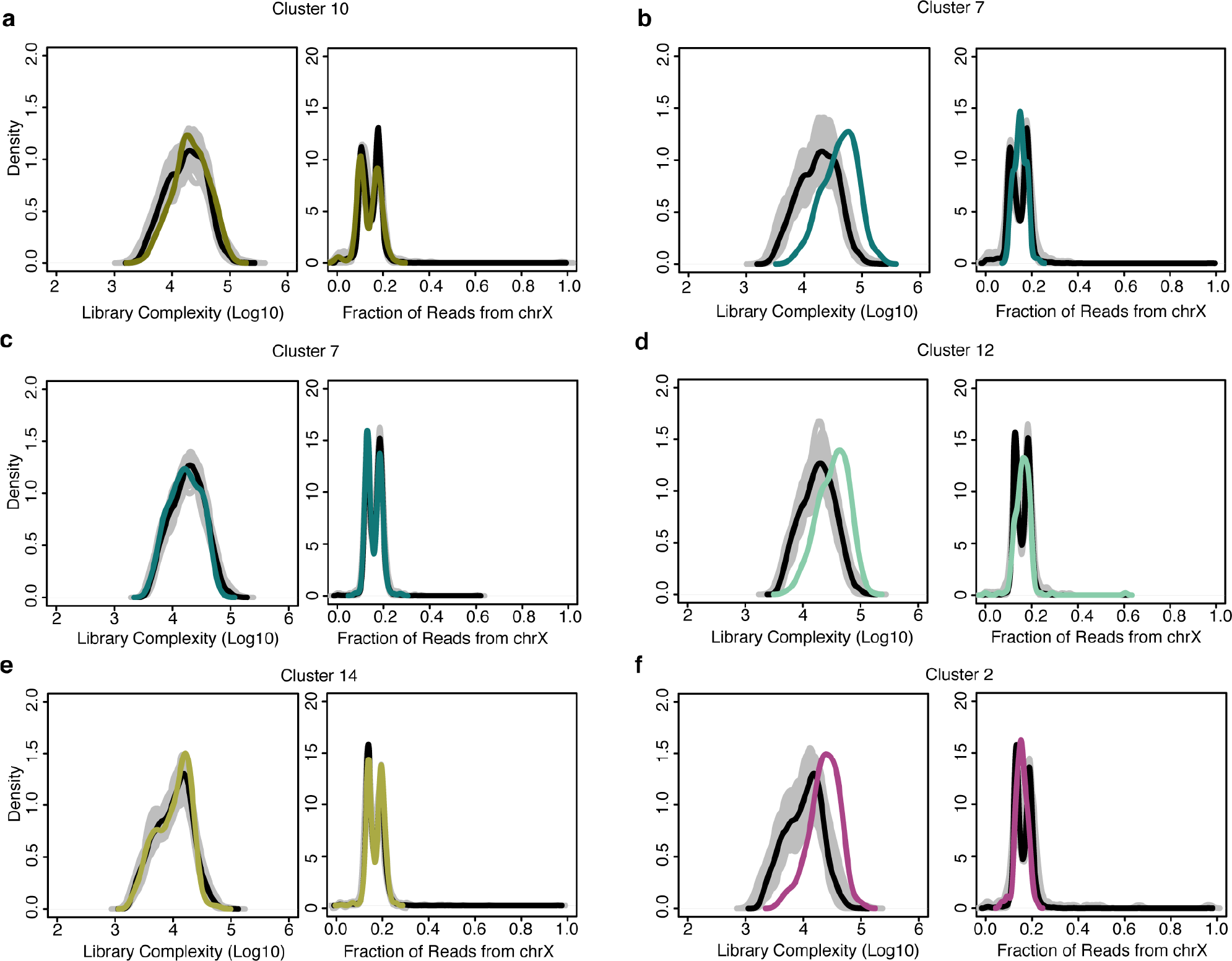
Library complexity and fraction of X chromosome reads highlights clusters of ‘collisions’ between cells from different tissues. Density plots of the estimated library complexity (using the same equation implemented in Picard; left panels) and the representation of chromosome X reads (right panels) in individual clusters. While most of the clusters defined by t-SNE are readily biologically interpretable, a small number of clusters (containing relatively few cells) defied easy characterization and are marked by an increases in both library complexity and an unusual distribution of X-chromosome to autosomal reads. These clusters likely correspond to clusters of “collisions” - cases in which two or more distinct cells share the same barcode as a consequence of the combinatorial indexing protocol. In each panel, the black line is the global distribution for all cells in that time point. The gray lines denote the results of randomly sampling an equal number of cells to the cluster in question. The colored line marks the distribution for the cluster being interrogated. **a**, **c**, **e**, Most clusters show relatively similar distributions of reads per cell (or library complexity, left) and a characteristic, bimodal distribution among cells in the ratio of X chromosome to autosomal reads (reflecting our use of a pool of male (XY) and female (XX) embryos, right). **b**, **d**, **f**, Putative collision clusters show a clear increase in the average library complexity (left) and a unimodal rather than bimodal distribution of X:autosomal reads (right). These features are not universally diagnostic (e.g. cluster 16 at 2-4 hrs does seem to show a strong, bona fide sex bias), but the combination of features is strongly predictive of clusters containing few cells and conflicting biological annotations based on gene/enhancer overlaps.

**Extended Data Figure 10.**
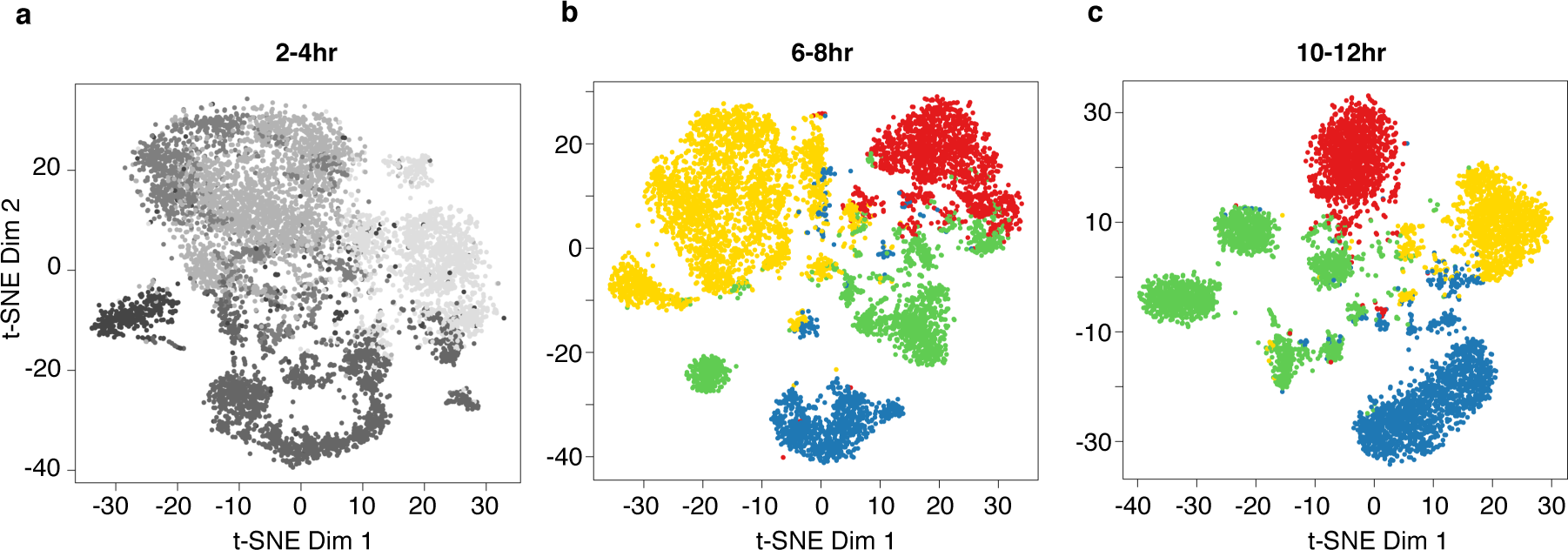
LSI defined clades and t-SNE clusters show strong correspondence. Shown are t-SNE clustered cells from each of the three time points colored by the LSI clade to which they were previously assigned (Fig. 1d-f) For the post-gastrulation time points, green is endoderm, red is myogenic mesoderm, yellow is non-neurogenic ectoderm, and blue is neurogenic ectoderm. There is strong correspondence between the germ layer level clade annotations from the LSI analysis with tissue-specific t-SNE clusters, particularly at the post-gastrulation time points (6-8 hrs and 10-12 hrs).

**Extended Data Figure 11.**
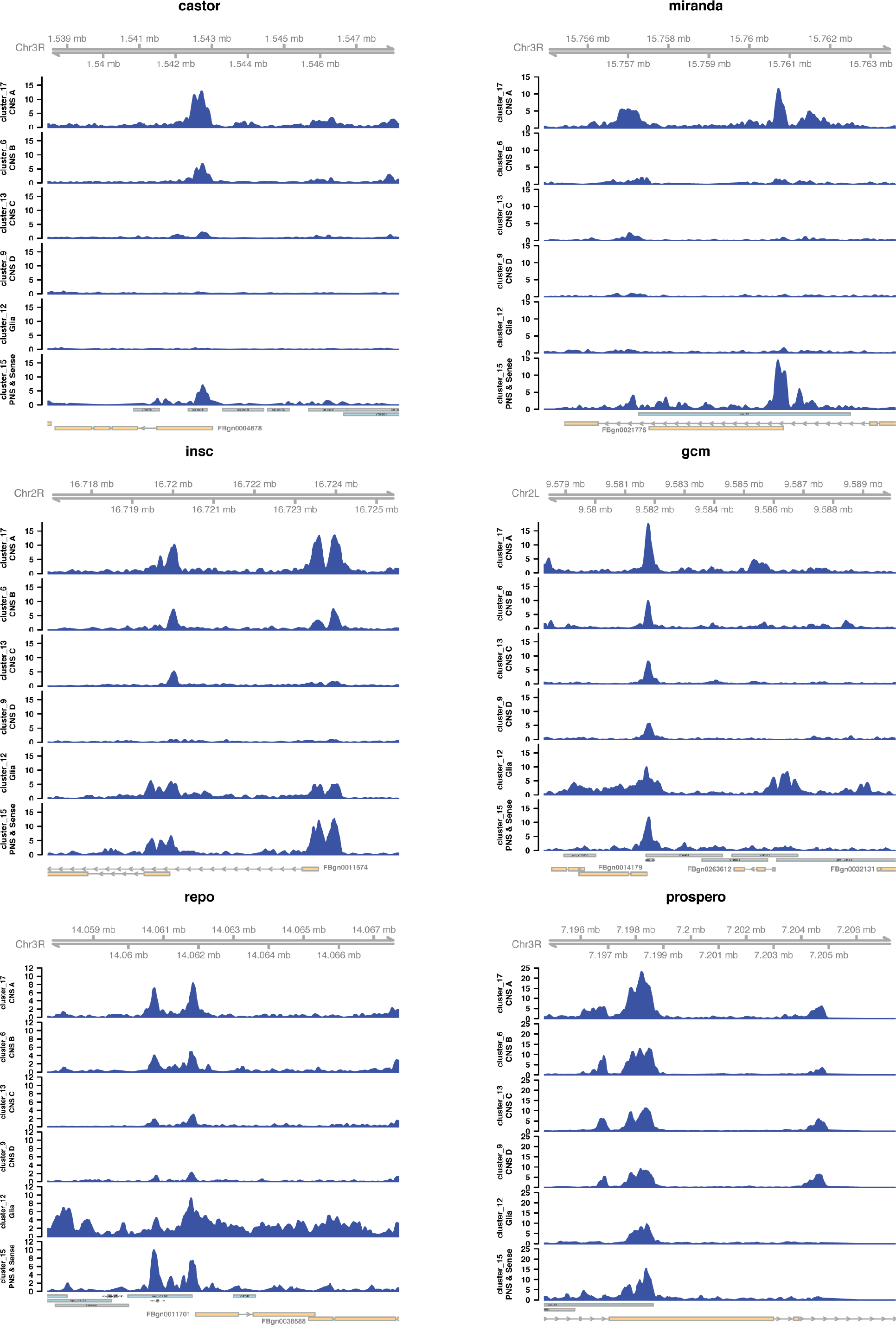
Differential accessibility around key regulators of neurogenesis between different t-SNE neural clusters. Clustering of 10-12 hr sci-ATAC-seq data using t-SNE reveals a wealth of neural related clusters. Of particular interest are a set of four distinct, but clearly related, clusters that are highly enriched for peaks overlapping enhancers active in the central nervous system (CNS). The identities of these clusters are not immediately clear, though they may represent the maturation process of neurons that is ongoing at 10-12 hours of development. Interestingly, the signatures of chromatin accessibility are highly different among these clusters in regions surrounding key regulators of neurogenesis (e.g. gcm, castor, or miranda). Shown here is normalized read coverage from neural clusters 17, 6, 13, 9 (annotated as CNS), cluster 12 (glia) and cluster 15 (PNS + sensory systems) around key drivers of neurogenesis at 10-12 hrs. Blue bars at bottom show the location of described transgenic reporters (known enhancers) from the CAD4 enhancer database (Table S11).

**Extended Data Figure 12.**
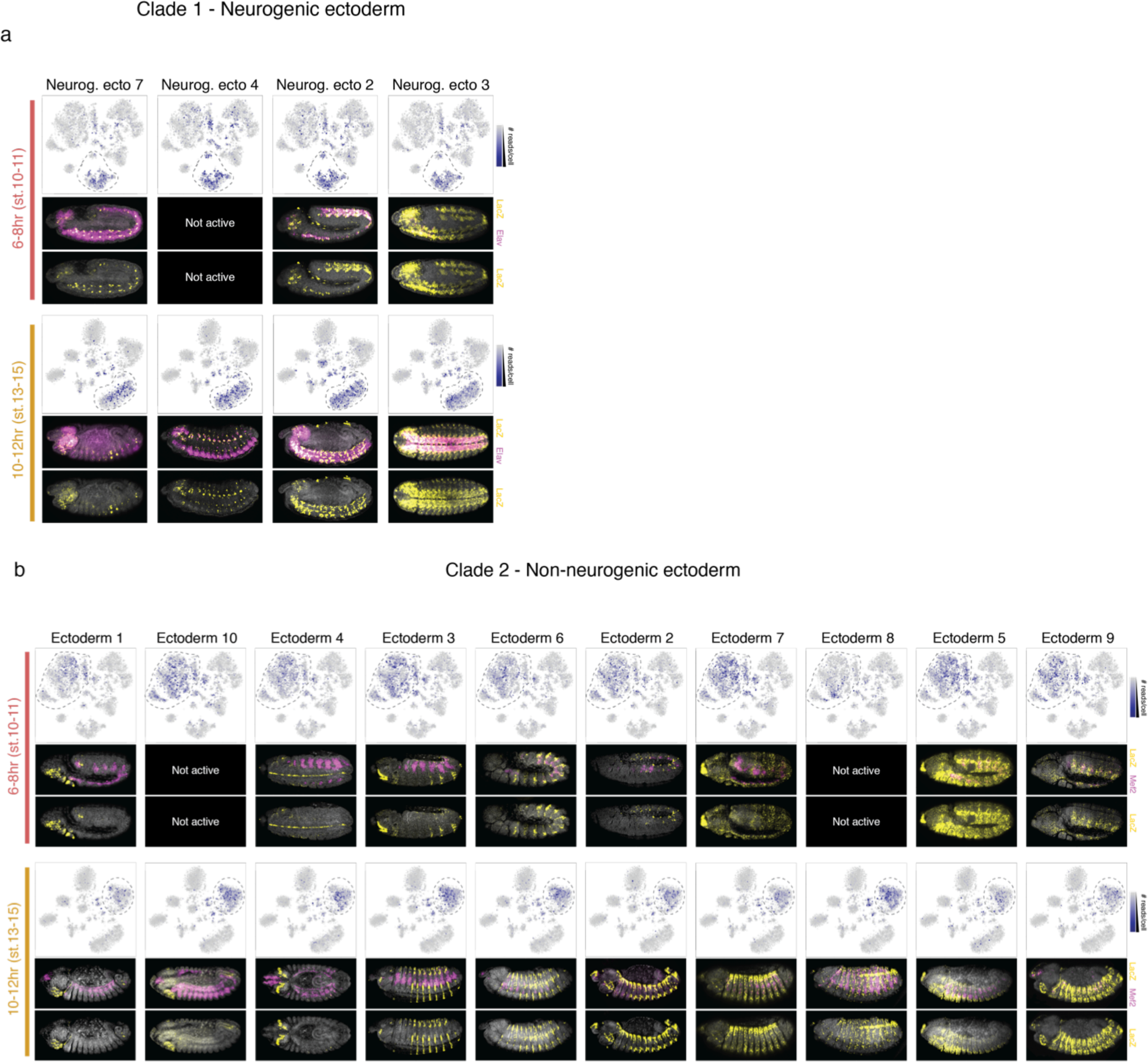
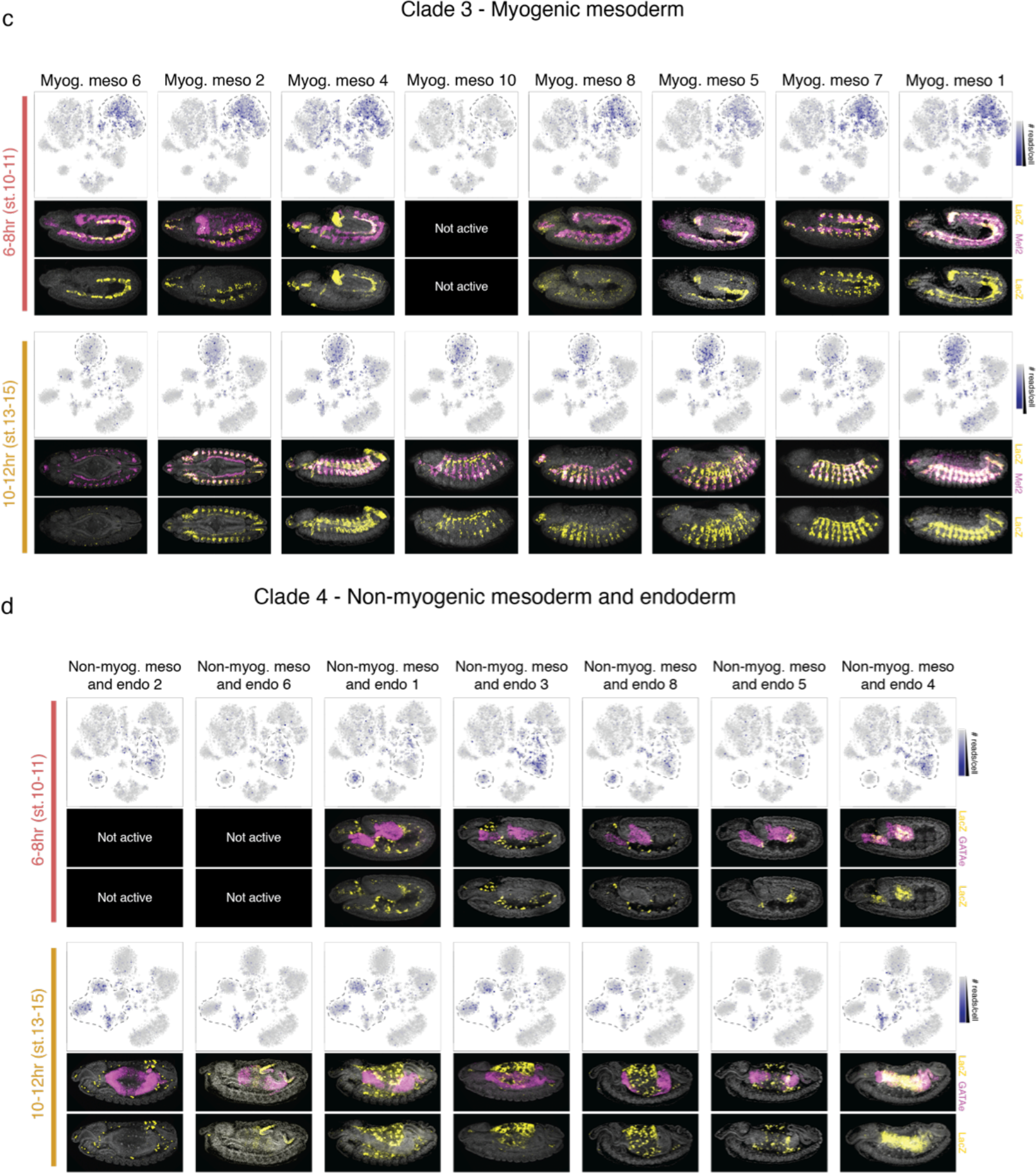
sci-ATAC-seq can predict tissue-specific enhancer usage during development. **a-d**, Candidate clade-specific enhancers tested in transgenic reporters: **a**, clade 1 (neurogenic ectoderm), **b**, clade 2 (non-neurogenic ectoderm), **c**, clade 3 (myogenic mesoderm), and **d**, clade 4 (non-myogenic mesoderm and endoderm). For each time point, upper panels show single cells visualized by t-SNE with the intensity of blue representing the number of sci-ATAC-seq reads obtained from each tested element in each individual cell. Cell clusters bounded by dashed lines correspond to the predicted clade of activity. Lower panels show representative embryos for each time point with nuclei stained with DAPI (grey), and in situ hybridization for the reporter gene driven by the enhancer (yellow) and a tissue marker (magenta).

**Table S1.** Summary of summits of accessibility. These are the identified regulatory elements (called in individual clades) used for downstream analyses. Clade assignment indicates the clade for which the peak was called specific. Clade 0 is reserved for peak calls that were not considered specific to any one clade at that time point.

**Table S2.** Enrichment analyses for intersections of distal accessible sites (>500 bp from any annotated transcription start site) for individual LSI-defined clades and a series of annotation datasets (see “Gene Expression, Enhancer Expression, and TF Binding Data” in Methods for details).

**Table S3.** Enrichment analyses for intersections of proximal accessible sites (within 500 bp of an annotated transcription start site) for individual LSI-defined clades and a series of annotation datasets (see “Gene Expression, Enhancer Expression, and TF Binding Data” in Methods for details).

**Table S4.** Enrichment analyses for intersections of all accessible sites for individual LSI-defined clades and a series of annotation datasets (see “Gene Expression, Enhancer Expression, and TF Binding Data” in Methods for details).

**Table S5.** Enriched motifs identified in clade-specific distal regulatory elements from the all three time points using SeqGL.

**Table S6.** Enrichment analyses for intersections of all accessible sites for individual t-SNE-defined clusters and a series of annotation datasets (see “Gene Expression, Enhancer Expression, and TF Binding Data” in Methods for details).

**Table S7.** Enrichment analyses for intersections of distal accessible sites (>500 bp from any annotated transcription start site) for individual t-SNE-defined clusters and a series of annotation datasets (see “Gene Expression, Enhancer Expression, and TF Binding Data” in Methods for details).

**Table S8.** Enrichment analyses for intersections of proximal accessible sites (within 500 bp of an annotated transcription start site) for individual t-SNE-defined clusters and a series of annotation datasets (see “Gene Expression, Enhancer Expression, and TF Binding Data” in Methods for details).

**Table S9.** Enrichment analyses for intersections of all accessible sites at the 2-4hr time point for individual t-SNE-defined clusters and a series of annotation datasets that were specifically compiled for this time point (see “Gene Expression, Enhancer Expression, and TF Binding Data” in Methods for details).

**Table S10.** Summary of candidate enhancers that were tested in transgenic assays and their observed activity in embryos. The sequences of the primers used to clone each region are also included.

**Table S11.** CAD4 database. A custom enhancer database of ~8,000 transgenic reporter assays covering 15% of the non-coding genome, including their tissue-specific activity.

## ONLINE METHODS

### Fixation of embryos and nuclear isolation

Wild-type *D. melanogaster* embryos were collected and fixed as previously described^33^. Briefly, embryos were collected on apple-agar plates in two-hour windows following three one-hour pre-collections to synchronize the collections. After aging (at 25°C) to the appropriate time window, embryos were washed from the plates, cleaned and dechorionated in 50% bleach for 2 minutes, followed by 15 minutes fixation while shaking at room temperature in a solution of heptane and cross-linking solution (1.8% formaldehyde v/v). Fixation was stopped by washing with a solution of 125 mM glycine in PBS. The embryos were then washed, dried, and frozen at -8°C in ~1 g aliquots. Embryo dissociation and nuclear isolation was obtained using a dounce homogenizer and a 22G needle (as described in steps 1-10^13^). The resulting nuclei were pelleted at 2,000g at 4C, resuspended in Nuclear Freezing Buffer (50 mM Tris at pH 8.0, 25% glycerol, 5 mM Mg(OAc)_2_, 0.1 mM EDTA, 5 mM DTT, 1X protease inhibitor cocktail [Roche], 1:2500 superasin [Ambion]), and flash frozen in liquid nitrogen.

### Collection of sci-ATAC-seq data

Our protocol for generating sci-ATAC-seq data was largely as previously described^1^, but with a few important improvements. Frozen nuclei were thawed quickly in a 37°C water bath and then pelleted at 500g for 5 minutes at 4°C, aspirated and resuspended in cold lysis buffer (supplemented with protease inhibitors). Nuclei were stained with 3 µM DAPI and 2,500 DAPI+ nuclei were sorted into each well of a 96-well plate containing 9 ul of lysis buffer (10 mM Tris-HCl, pH 7.4, 10 mM NaCl, 3 mM MgCl2, 0.1% IGEPAL CA-630 from^34^ supplemented with protease inhibitor (Sigma)) and 10 µl of TD buffer (Illumina, part of FC-121-1031) in each well. 1 µl of each of the 96 custom and uniquely indexed Tn5 Transposomes (Illumina, 2.5 uM)^35^ was then added to each well and nuclei were incubated at 55°C for 30 minutes. Following tagmentation, 20 µl of 40 mM EDTA (supplemented with 1 mM Spermidine) was added to stop the reaction and the plate was incubated at 37°C for 15 minutes. All wells of the plate were then pooled, nuclei were stained again with 3 µM DAPI and 25 DAPI+ nuclei were sorted into each well of a second set of 96-well plates that contained 12 µl of our reverse crosslinking buffer (EB buffer [Qiagen] supplemented with 833 µg/ml Proteinase K [Qiagen] and 0.04167% SDS). For each time point, we collected 4 plates of nuclei at this stage. We expect that sorting 25 nuclei into each well at this stage will result in approximately 12% of barcodes representing more than one nucleus^1^. Nuclei were then incubated overnight at 65°C. Proceeding from reverse-crosslinking, we added primers (0.5 µM final concentration), 7.5 µl of NPM polymerase master mix (Illumina, FC-121-1012) and BSA (2X final concentration; NEB) to each well. Tagmented DNA was then PCR amplified. To determine the number of cycles required, we first amplified several test wells of nuclei that had been sorted onto an additional plate and monitored the reactions with SYBR green on a qPCR machine to establish when the libraries reached saturation. The cycling conditions were as follows:
72°C 3 minutes
98°C 30 seconds
15-25 Cycles:
98°C 10 seconds
63°C 30 seconds
72°C 1 minute
Hold at 10°C

We have found that the optimal number of cycles can vary from one experiment to the next, but is usually in the range of 15-25 cycles. After PCR amplification, all wells were pooled and then split across 4 DNA Clean & Concentrator-5 columns (Zymo) and then all 4 products were pooled and cleaned again using Ampure beads (Agencourt). Finally, the concentration and quality of the libraries was determined using the BioAnalyzer 7500 DNA kit (Agilent). For sequencing, equimolar libraries from the three time points were pooled and loaded 1.5 pM on a NextSeq High output 300 cycle kit and sequenced using custom primers and a custom sequencing recipe. 50 base pairs (bp) were sequenced from each end, as were the barcodes introduced during tagmentation and PCR amplification.

### Read alignment, cell assignment, and duplicate removal

To process the data, BCL files were converted to fastq files using bcl2fastq v2.16 (Illumina). Each read was assigned a barcode which was actually made up of 4 individual components: a tagmentation barcode and a PCR barcode added to the P5 end of the molecule and a distinct tagmentation and PCR barcode added to the P7 end of the molecule. To correct for sequencing or PCR amplification errors, we broke the barcode into its constituent parts and matched each piece against all possible barcodes. If the component was within 3 edits of an expected barcode and the next best matching barcode was at least 2 edits further away, we fixed the barcode to its presumptive match. Otherwise, we classified the barcode as ambiguous or unknown. We next mapped each read to the dm3 reference genome using BWA aln and then filtered out read pairs that did not map uniquely to autosomes or sex chromosomes with a mapping quality of at least 10, as well as reads that were associated with ambiguous or unknown barcodes. We subsequently removed all duplicate reads using a custom python script that only considered reads assigned to the same barcode. Finally, to determine which barcodes represented genuine cells (as opposed to background reads assigned to improper barcodes), we counted the number of reads assigned to each barcode and used the mclust package in R^36,37^, which fits the data using a mixture model and determines the maximum likelihood parameters for a given number of distributions, to define two distributions of barcodes – setting the read depth cutoff for a cell at the point at which we were 95% confident that the barcode belonged to the higher read depth distribution. Considering the distribution of barcodes for all three experiments at the same time, we determined this read depth cutoff to be 518 reads (i.e. we required a barcode to be associated with at least 518 reads to be considered a true cell; Extended Data Fig. 1).

### Latent semantic indexing

To further process the raw data we first broke the genome into 2kb windows and then scored whether each cell had any insertions in each window, creating a large binary matrix of windows by cells for each time point. Based on this binary matrix, we filtered out the lowest 10% of cells in terms of number of sites accessible and then only retained the top 20,000 most commonly used sites (this number could extend a little above 20,000 because we retained all sites that were tied at the threshold for cell counts). We then normalized and re-scaled these large binary matrices by using the term frequency-inverse document frequency (“TF-IDF”) transformation. We first weighted each site that was accessible in an individual cell by the total number of sites accessible in that cell. We then multiplied these weighted values by log(1 + the inverse frequency of each site across all cells). Subsequently, we performed singular value decomposition on the TF-IDF matrix and then generated a lower dimensional representation of the data by only considering the 2nd through 6th dimensions (because we have found that the first dimension is always highly correlated with read depth). These LSI scores of accessibility were then used to cluster cells and windows based on cosine distances using the ward algorithm in R. Visual examination of the resulting bi-clustered heatmap identified 4-5 major clades for each time point.

### Peak calling

In order to identify specific regulatory elements within each of the major clades in each time point, we aggregated the data across cells from each clade using a process we call “*in silico* cell sorting”. To do so we simply collected all the unique mapped reads associated with cells that were assigned to a given clade and saved that as a distinct bam file. Then for each bam file representing a clade, we used the MACS2 program^26^ to identify peaks of increased insertion frequency, as well as summits of accessibility within each of those peaks. For MACS, we used the macs2 callpeak command with the following parameters: –nomodel –keep-dup all –extsize 200 –shift -100. For downstream analyses we generated a master list of potential regulatory elements by taking 150 bp windows centered on all summits called in each clade in each time point and merged them with the BEDTools program^38^. For Fig. 1c, we also compared our sci-ATAC-seq data to DNase-seq data collected by Thomas et al^3^ on whole embryos at similar time points. In order to be consistent in our comparisons (and provide a comprehensive list of peaks), we downloaded the raw DNase-seq reads (36 bp, single-end), remapped them with our pipeline and called peaks with MACS2 as described above. Specifically, we downloaded two replicates for each of 3 time points - stage 5, stage 11 and stage 14. Peaks called on each replicate independently were intersected to create a master list of peaks for each time point. These peak lists were then intersected with our sci-ATAC-seq data.

### Identification of differentially accessible sites

To identify regulatory elements that were more accessible specifically in individual clades, we started by generating a new binary matrix of insertion scores for individual cells using the master list of summits of accessibility described above. We could then use a logistic regression framework to test whether cells of a given clade were more likely to have insertions at a given site relative to all other cells. To identify sites that were specifically more accessible in a single clade, we first found summits that were significantly more open in a given clade at a 1% false discovery rate (FDR). To ensure that these sites were specific to any one clade, we also filtered out sites that were significantly open in any other clade at a relaxed 20% FDR. All testing of differential accessibility was implemented with the Monocle package^39^ using the binomialff test. For this analysis, only sites observed in at least 50 cells in a given time point were tested.

### K-mer discovery

We used the SeqGL program^40^ to identify motifs that were enriched in clade-specific elements. To do so, we started with all sites were specifically more accessible in the clade based on our logistic regression testing described above. Because our master list of sites included sites of variable length (after merging all sites from all clusters), we only considered 150 bp windows centered on summit midpoints. We also filtered out any sites within 500 bp of a transcription start site (to focus on the more tissue specific distal elements). As a background set of regions we randomly selected an equal number of sites from the master summit list that matched the GC content and repeat element content of the test set (this was controlled using a script provided in the gkm-SVM software package)^41^. Finally, instead of default parameters, we used 200 groups and 30,000 features, similar to the parameters used to analyze DNase-seq data in the original SeqGL publication.

### Gene expression, enhancer expression, and TF binding data

To perform categorical enrichments, we annotated regions/windows/peaks of the non-coding genome using two types of experimental information: 1) tissue specific expression of the nearest gene, from *in situ* hybridization data and 2) a custom enhancer database of ~8,000 transgenic reporter assays covering 15% of the non-coding genome (CAD4; Table S11). *Drosophila*-specific gene-level functional information (biological process, molecular function, and cellular compartment) was downloaded from the Gene Ontology Consortium (v.1.2). Additional, higher-level functional annotations were downloaded from the PANTHER classification system (v.8) corresponding roughly to the higher-level categories of the GO-SLIM ontology. Gene expression information was obtained from the Berkeley *Drosophila* Genome Project *in situ* database (http://insitu.fruitfly.org/cgi-bin/ex/insitu.pl) and from a download of the FlyBase gene expression annotations (May 2016). Almost all expression terms from these two sources could be mapped to a common set of hierarchically organized anatomical terms (FlyBase anatomy OBO file v.1.47). In the few cases where an exact match could not be found, a choice was made manually or using the map provided by FlyBase (FBrf0219073). The stage/timing information from both datasets was shifted as needed to match a common set of grouped stages (stages 1-3, stages 4-6, stages 7-8, stages 9-10, stages 11-12, stages 13-16). Our compiled data is available in Table S11.

To more directly assay the regulatory activity of the non-coding genome, we made use of an updated version of the CRM Activity Database (CAD4; Table S11). This dataset consists of a combination of three primary resources: Our previous CAD enhancer database^42^, entries from the RedFly enhancer database (Release 5)^6^, and data from the Vienna Tiling Project^5^. We compiled this dataset in two steps. First, all expression terms (and timing terms, where available) were mapped to a common standard (FlyBase anatomy terms v.1.47) and, when timing information was available) a common set of stage windows (stages 1-3, stages 4-6, stages 7-8, stages 9-10, stages 11-12, stages 13-16). In most cases, the mapping was automatic and unambiguous. In a few cases, manual term matching was required (generally unambiguous). In the second step, we merged overlapping entries from CAD3 and the RedFly database and manually removed redundant information. Given the different methodologies used in the compilation of the data sources, no attempt was made to combine entries from CAD3/RedFly with the Vienna Tiles.

To further explore the function of specific regions of noncoding DNA, we also made use of a custom compilation of high-quality transcription factor binding data from ChIP studies during embryogenesis (taken from ref^32^) that allowed us to assign transcription factor binding events to each window/peak. Transcription factor binding motifs were taken from this same dataset. To infer likely transcription factor binding events, we scanned under published ChIP peaks for instances of the motif using FIMO^43^.

### Categorical enrichments

To identify enriched categories within our LSI clades, we first assigned categorical labels by looking for overlaps between our enhancer expression database and our summit regions, with summit regions inheriting the timing and expression labels of all overlapping elements. Gene-based annotations (expression, GO, and PANTHER terms) were assigned by association to the nearest gene.

To identify differentially accessible summit regions we used a logistic-regression framework (see above) as applied to all summit regions containing reads in at least 50 cells. Enriched summit regions constituted the foreground set for any clade, with the remaining tested summit regions constituting the background set. For each of our category sets (e.g. enhancer expression, gene expression, or GO) we used a Fisher’s exact test to look for over-representation of each category among our foreground set relative to the background set. Because many of our categories are strongly overlapping, we have applied no formal correction for multiple comparison, choosing instead to focus on large, consistent enrichments with highly significant p-values. Overlaps among significant categories were visualized by plotting distances between categories using the pyEnrichment package (https://github.com/ofedrigo/pyEnrichment) to avoid overcalling a category.

Categorical enrichment within our t-SNE clusters was assessed similarly. Foreground sets per cluster (within each time point) were assessed using the results of our binomial enrichment test (q-value <= 0.01 and a beta > 0). The background set consisted of all other tested summits at that time point (see above).

### t-stochastic neighbor embedding and cluster identification

To identify clusters of cells with finer resolution than the LSI-based clades, we used t-SNE^18^ for dimensionality reduction. To do so, we started with the same binary matrix of insertions in summits that we used to identify clade-specific differentially accessible sites. We again filtered out the lowest 10% of cells (in terms of site coverage) and in this case we only retained sites that were observed in at least 5% of cells. We then transformed this matrix with the TF-IDF algorithm described above. Finally, we generated a lower dimensional representation of the data by including the first 50 dimensions of the singular value decomposition of this TF-IDF-transformed matrix. This representation was then used as input for the Rtsne package in R^18,27,28^. To identify clusters of cells in this two dimensional representation of the data, we used the density peak clustering algorithm^44^ as implemented in Monocle 2^29^. Rho and delta parameters were chosen to be as inclusive of outlier peak centers as possible (based on the decision plot), while making sure that the clusters were sensible based on visual inspection of the cluster assignments on the t-SNE plot.

### t-SNE differential accessibility

To identify summits that were significantly more accessible in t-SNE-defined cell clusters, we used a similar framework to the one described for LSI-based clades above. There were however a few differences. In this case, we consider sites that were seen in at least 10 cells in any time point (instead of 50). In addition, we did not use a second cutoff to determine specificity within a time point.

### Sexing individual nuclei

Another biological axis of the data that came to light through the use of t-SNE plots was that we were able to clearly distinguish nuclei from male and female embryos. In an initial analysis, we included data from the sex chromosomes while clustering cells (as was done for the germ layer analysis). This resulted in many individual cell clusters appearing ‘bi-lobed’ (Extended Data Fig. 7a), which prompted us to explore if there was sex-bias in the lobes of individual cell clusters. We found that the distribution of reads mapping to the X chromosome in individual cells was distinctly bimodal (Extended Data Fig. 7b), allowing us to assign a sex to each cell. When we colored the t-SNE plots by these sex assignments we found that the lobes of individual cell clusters almost perfectly segregated the sexes (Extended Data Fig. 7c). To alleviate this problem we excluded sex chromosome reads from our analysis and re-clustered cells with t-SNE. This resolved the ‘bi-lobed’ structures and removed the sex bias from almost every individual cluster (Extended Data Fig. 7d).

### Identifying clusters of cells that are likely artifacts of barcode collisions

Several small clusters (e.g. cluster 12 at 6-8 hrs) appear to be mixtures of cells from different germ layers and/or tissues based on our enrichment analysis. To determine if these are technical (due to barcode collisions, where one cell barcode represents the nuclear contents of two cells) or biological we used two metrics to identify collisions (instances wherein two or more cells coincidentally pass through the same combination of wells during sci-ATAC-seq): First, we looked at the estimated complexity of individual cells that make up these small clusters, as collisions are expected to be twice as complex on average as barcodes that truly represent an individual cell. Second, we considered whether the proportion of reads mapping to the X chromosome for cells in these clusters was distinctly bimodal, as collisions would be just as likely to combine data from cells of the opposite sex as from two cells of the same sex (Extended Data Fig. 9). While the vast majority of clusters exhibited distributions of complexity and X chromosome coverage consistent with single nuclei, a small subset of clusters in each time point showed either higher complexity than expected, more unimodality of reads mapping to the X chromosome, or both - consistent with our suspicion that these are cell collision clusters (Extended Data Fig. 9). At 2-4 hrs, we identified one, at 6-8 hrs three and 10-12 hrs five potential collision clusters (Fig. 4a,b purple clusters).

### Transgenic enhancer assays

Candidate clade-specific enhancers were selected from sci-ATAC-seq summits using the following criteria only: (1) Summit shows enriched accessibility specifically in the target cell clade at 6-8 hrs and/or 10-12 hrs (q-value<0.01 and beta>0 in target clade, q-value>0.2 in all other clades); (2) summit does not fall within 500 bp of an annotated transcription start site; (3) summit does not overlap a region already in our enhancer database. Summits showing a range of effect sizes (beta) were selected (minimum beta approx. 1.9; see Table S10). The selected regions, plus 100-200 bp of flanking sequence, were PCR amplified from genomic DNA (primers are listed in Table S10) and cloned upstream of a minimal *hsp70-*promoter driving a *LacZ* reporter gene in an attB-containing plasmid. All constructs were injected into embryos according to standard methods^45^ and inserted into the attP landing site line M{3xP3-RFP.attP’}ZH-51C via PhiC31 integrase insertion^46^, yielding integration at chromosomal position 51C1. Transgenic lines were generated by BestGene Inc (Chino Hills, CA, USA). Overnight collections of homozygous embryos spanning all stages of embryogenesis were formaldehyde-fixed, stained by double fluorescent *in situ* hybridization^47^, and mounted in ProLong Gold with DAPI (Invitrogen; cat. #P36931). Antisense *in situ* probes against *LacZ* and a tissue marker gene were used: *Mef2* marking myogenic mesoderm was used for predicted myogenic mesoderm and non-neurogenic ectoderm enhancers; *GATAe* was used for predicted non-myogenic mesoderm and endoderm enhancers. For the predicted neurogenic ectoderm enhancers, neurons were marked by immunostaining with antibodies against the Elav protein (Elav-9F8A9; Developmental Studies Hybridoma Bank). Images were acquired with a Zeiss LSM780 laser-scanning confocal microscope using a PlanApo 20X/NA0.8 objective at an effective pixel size of 461 nm in xy. Images were processed using Fiji^48^. Annotated t-SNE plots for each candidate enhancer were produced by plotting the sum of sci-ATAC-seq reads per cell that overlap each tested genomic region.

